# Transcriptomic signatures of mouse ovarian aging and estropausal transition at single cell resolution

**DOI:** 10.1101/2024.08.12.607592

**Authors:** Xifan Wang, Jiping Yang, Chen Jin, Xizhe Wang, Daniela Contreras, Melody Devos, Yousin Suh

**Affiliations:** Department of Obstetrics and Gynecology, Columbia University, New York, NY10032, USA

## Abstract

Female reproductive aging affects fertility and overall health. Functional decline in mouse ovary during reproductive aging is accompanied by estrous cycle prolongation and cessation. However, the molecular mechanism underlying reproductive aging concomitant with such cycle changes remains unclear. Using single-cell transcriptomics, we characterized aging signatures in mouse ovaries across the reproductive lifespan ranging from reproductively young (regular cycle) through peri-estropause (regular vs. irregular cycles) to post-estropause age (acyclic). Reproductive aging significantly remodeled cell compositions and increased transcriptional heterogeneity, with more pronounced changes post-estropause, exhibiting coordinated alterations across cell types in the ovary. Genes undergoing monotonic changes during reproductive aging across cell types were consistently enriched in the conserved pathways of aging, including oxidative phosphorylation, stress responses, and proteostasis. Additionally, cell type-specific changes were identified including dysregulation of hormone synthesis in granulosa cells, alterations in collagen and hyaluronan metabolism in stromal and early theca cells, and functional decline of a unique phagocytosis-associated macrophage. Aging also led to a significant decrease in cell-cell communications, particularly between stromal and granulosa cells, and an increase in extracellular vesicle secretion. Furthermore, we found increased expression of the senescence marker Cdkn1a and senescence-associated secretory phenotype (SASP) factors during ovarian aging, especially in granulosa cells. Notably, most of these aging-associated changes were more pronounced in irregular cycling ovaries compared to the regular cycling counterparts at the same age during the peri-estropause stage, suggesting that aging-related molecular changes in the ovary drive the estropausal transition in mice.

## Introduction

The ovary is the key organ that maintains female reproductive and endocrine function, undergoing accelerated aging compared to other organs^1^. Ovarian aging refers to the progressive decline of ovarian functions with age, characterized by the decrease in the quantity and quality of oocytes residing within the follicles^2^. The gradual loss of ovarian function not only contributes to infertility and endocrine dysfunction but also leads to deleterious consequences for overall health^3,4^.

Mice share a similar process of ovarian aging with women, which can be broadly divided into three phases as reproductively young, the estropausal transition (akin to perimenopausal in women), and post-estropause (similar to post-menopause in women)^5–7^. Mice have regular estrous cycles occurring every four to five days at reproductively young age, consisting of four phases, proestrus, estrus, metestrus, and diestrus^8^. The functional decline in mouse ovaries is accompanied by estrous cycle prolongation and cessation. Around 9–12 months of age, mice enter the estropausal transition stage, during which they may experience either regular or irregular estrous cycles^6^. The cessation of cyclicity (acyclicity) typically occurs between 13 and 16 months of age, termed as post-estropause^9^. However, our understanding of basic molecular mechanisms underlying mouse ovarian aging across the entire reproductive lifespan concomitant with such cycle changes remains limited. Furthermore, there is considerable heterogeneity in the onset of cycle irregularity in mice, even with identical genetic backgrounds and environments, similar to the inter-individual variability in the age of peri-menopausal transition observed in women^10^. The molecular mechanisms driving the onset of cycle irregularity at the estropausal transition remain unclear.

The ovary encompasses various cell types including oocytes surrounded by granulosa and theca cells within follicles, alongside supporting somatic cells like stromal, vascular, and immune cells, collectively orchestrating its dynamic functions in follicular development and hormone production. During follicle development and ovulation, granulosa cells (GCs) proliferate and form into different layers, while surrounding stromal cells are recruited and promoted to differentiate into specialized theca cells (TCs)^11^. Meanwhile, the TC layers are infiltrated by immune cells, which play a vital role in inflammation and ovarian physiological processes such as follicle atresia, ovulation, corpus luteum formation and regression^12^. Macrophages, the most abundant immune cells with high levels of heterogeneity and plasticity in the ovary, are the central components of the inflammatory immune response and perform multiple functions to maintain tissue homeostasis^13^. Ovarian aging is a complicated process which is associated with the reduced ability of granulosa cells to counteract reactive oxygen species^14^, increased fibrosis in stromal extracellular matrix molecules^15^, and aberrant immune microenvironment^16^, in addition to the quality and quantity decline of oocytes. Due to the high heterogeneity of ovarian cell types and the complexity of cell-cell interaction, single-cell RNA sequencing (scRNA-seq) is invaluable for exploring the differentiation trajectory of ovarian somatic cells such as GCs and TCs, revealing cell type-specific alterations in gene expression, and unraveling changes in cellular communication during ovarian aging, especially for rare cell types. A recent study comparing single-cell transcriptomics of ovaries between 3-month and 9-month mice, unveiled an increase in lymphocytes and stromal fibrosis as early markers of ovarian decline^17^. Moreover, the impact of estrous cycling and aging on all female reproductive tract organs in mice at single-cell resolution was investigated and it demonstrated that the increased fibroblast inflammation and fibrosis played a central role in the aging of the uterus and oviduct, but showed minimal impact on the ovary^18^. Nevertheless, the impact of aging and cycle transition on subtypes of mouse ovarian cells, especially granulosa cells, stromal cells, theca cells, and macrophages, requires further investigation.

In this study, we systematically characterized the temporal dynamic signature of ovarian aging by performing scRNA-seq of mouse ovaries with a defined estrous cycle stage across the reproductive lifespan. Furthermore, we unraveled the cellular and molecular alteration in mouse ovaries associated with the onset of cycle irregularity by comparing irregular and regular cycling ovaries during the estropausal transition stage. Since it has been recently shown that there are dynamic transcriptional changes between different phases of the estrous cycle in young mouse ovaries^19^, we standardize the cycle phase of mice (diestrous) from different ages to mitigate confounding factors arising from cycling itself. This study provides an in-depth understanding of cell type-specific mechanisms of mouse ovarian aging at single-cell and temporal resolution concomitant with cycle changes. Our findings enabled the discovery of reproductive aging-associated biomarkers and the identification of cellular and molecular programs that can be potentially targeted to improve aging-associated ovarian disorders.

### Comprehensive cellular and molecular taxonomy of mouse ovary based on scRNA-seq

To comprehensively dissect the temporal dynamic signature of mouse ovarian aging, we conducted single-cell RNA sequencing (scRNA-seq) of mouse ovaries from reproductive young (Y, 4.5-month, n=5), peri-estropause (M, 10.5-month, n=7), and post-estropause (O, 15.5-month, n=5) stage **(Fig. 1a)**. The estrus cycle of each mouse was assessed daily for a minimum of 3 weeks. The reproductive young mice, exhibiting regular cycles every 4-5 days for at least 2 weeks, and the post-estropausal mice reaching acyclicity (no cycle lasting more than 9 days), were selected for ovarian scRNA-seq **(Fig. S1a)**. Among the peri-estropausal mice, 3 out of 7 displayed regular cycles, while the remaining 4 showed irregular cycles (2 contiguous cycles of 5–8 days, **Fig. S1a)**. All ovary samples were collected when the mouse was in diestrus.

**Figure 1.**
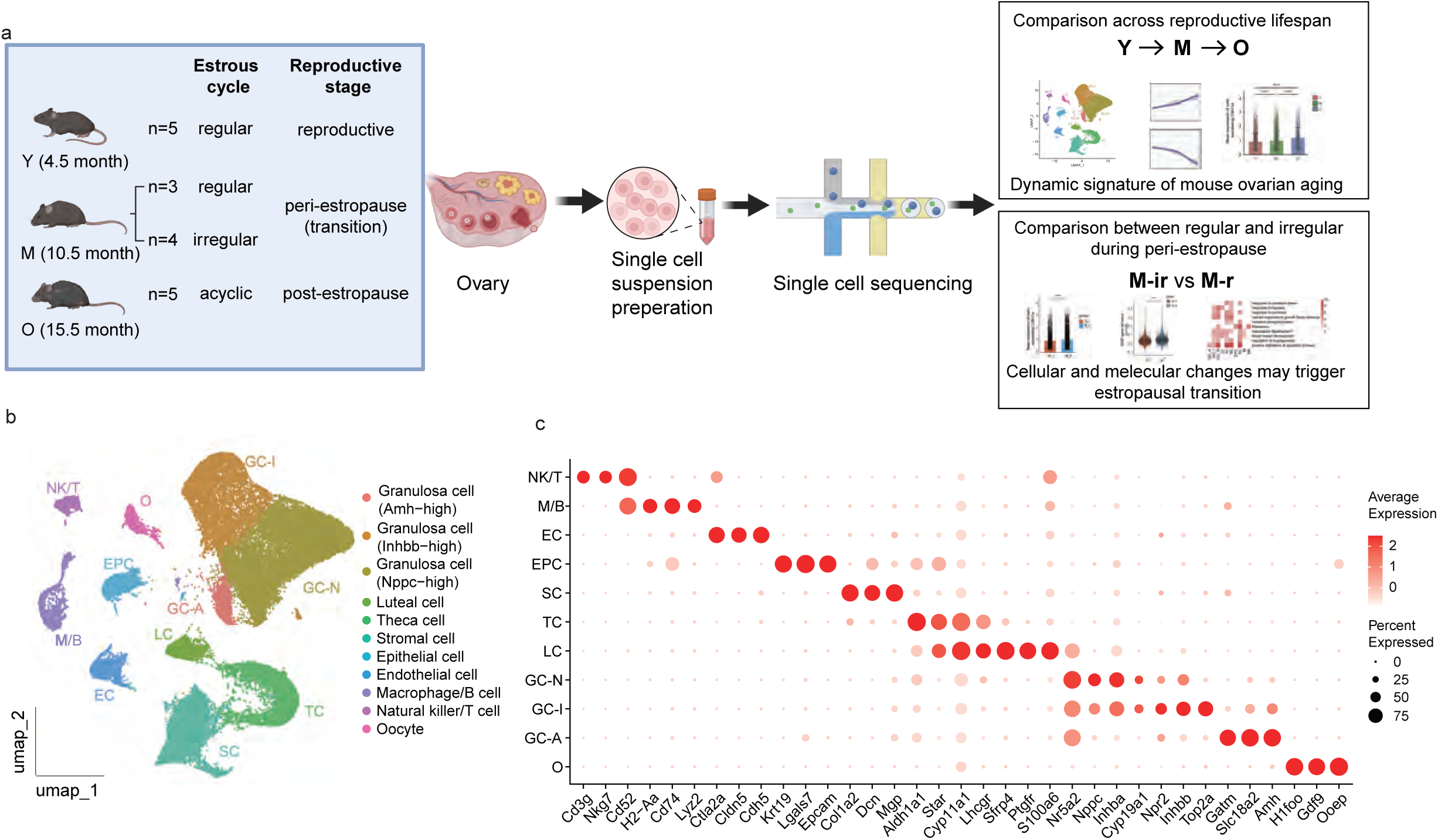
scRNA-Seq profiling of mouse ovaries. **a,** Flowchart overview of single-cell RNA-seq of mouse ovaries from different reproductive stages. **b,** Uniform manifold approximation and projection (UMAP) plots showing the cell types of mouse ovary. O, oocyte; GC-A, Amh-high granulosa cell; GC-I, Inhbb-high granulosa cell; GC-N, Nppc-high granulosa cell; LC, luteal cell; TC, theca cell; SC, stomal cell; EPC, epithelial cell; EC, endothelial cell; M/B, macrophage/B cell; NK/T, natural killer/T cell. **c,** Dot plots showing the expression of representative genes for each cell type.

After strict quality control (see Methods), we obtained 53,119 single-cell transcriptomes for downstream analysis and applied uniform manifold approximation and projection (UMAP) analysis to resolve the cell type distribution for each time point **(Fig. 1b and Fig. S1b)**. 11 distinct cell clusters were identified through unsupervised clustering. Based on well-defined cell type-specific markers, the major cell categories of the ovary were annotated, including oocytes (O, *Ooep*^+^), 3 groups of granulosa cells (GC), luteal cell (LC, *Sfrp4*^+^), theca cells (TC, *Aldh1a1*^+^), stromal cells (SC, *Dcn*^+^), epithelial cells (EPC, *Epcam*^+^), endothelial cells (EC, *Cdh5*^+^) and 2 groups of immune cells (IC) **(Fig. 1b and 1c)**. Granulosa cells encompassed three clusters: Amh-high granulosa cell (GC-A), Inhbb-high granulosa cell (GC-I), and Nppc-high granulosa cell (GC-N). Granulosa cells secret hormones such as Anti-Müllerian hormone (AMH), inhibin, and estradiol at different stages of folliculogenesis^20^. Interestingly, GC-A, GC-I, and GC-N exhibited a flow on the expression level of several hormone-related genes, including *Amh*, *Inhbb* and *Cyp19a1* et.al. This pattern suggests that these three GC clusters may represent granulosa cells from different stages of follicular development **(Fig. S1c)**. For instance, GC-A exhibited the highest expression of *Amh*, along with high expression of *Gatm*, indicating that this cluster may originate from pre-antral follicles. GC-I showed the highest expression of *Inhbb*, coupled with high expression of *Npr2* (a marker of cumulus cells) and Top2a (a proliferation marker), suggesting that this cluster likely originates from early-antral or antral follicles. GC-N demonstrated the highest expression of *Inhba* and *Cyp19a1*, in addition to high expression of *Nppc* and *Mro*, which are markers of mural granulosa cells in antral and preovulatory follicles. Both theca cells and luteal cells showed high expression of steroidogenic acute regulatory protein gene *Star*, luteinizing hormone/choriogonadotropin receptor gene *Lhcgr*, and progesterone synthesis enzyme *Cyp11a1*, indicating their contribution to progesterone generation. Notably, all clusters of granulosa cells and luteal cells showed expression of *Nr5a2* **(Fig.1c)**, a crucial transcriptional regulator modulating granulosa cell-specific gene regulatory network^21^, suggesting that *Nr5a2* could be a common marker for granulosa cells, including the luteinized granulosa cells. Immune cells contained two distinct clusters: one exhibited high expression of *Cd3g* and *Nkg7*, indicative of the natural killer cell and T lymphocyte cluster; the other was characterized by high expression of *Lyz2* and *Cd74*, representing the macrophage and B lymphocytes cluster.

### Altered cell distribution in mouse ovary during reproductive aging

To investigate the temporal change of cell distribution in mouse ovaries across the reproductive lifespan, we compared the proportions of different cell types in ovaries among reproductive young (Y), peri-estropause (M), and post-estropause (O) groups. Age-dependent changes in cell proportion of several cell types were observed **(Fig. 2a)**. The proportion of oocytes dramatically decreased in middle age when compared with the young ovaries and was nearly depleted in the old ovaries **(Fig. 2a)**. Similarly, the proportion of Amh-high granulosa cells, representing granulosa cells from early-stage follicles, gradually declines with aging and shows a significant difference in the old ovaries compared to the young/middle-aged ovaries **(Fig. 2a)**. In addition, theca cell abundance significantly decreased since middle age **(Fig. 2a)**. These changes could be attributed to the decline of follicle numbers during aging, particularly the number of small follicles, which is an important indicator of ovarian reserve.

**Figure 2.**
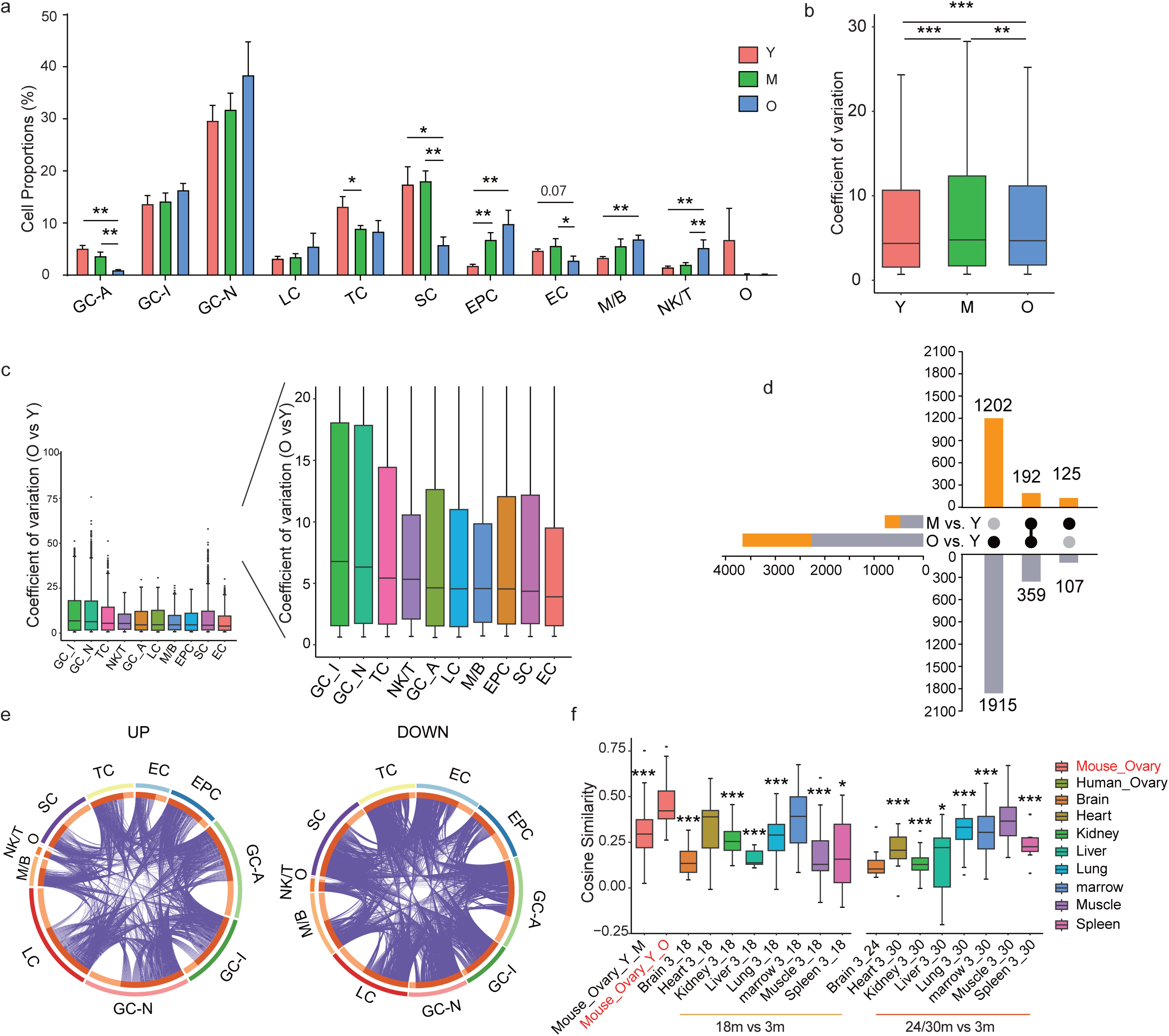
Cell distribution and global transcriptional change during mouse ovarian aging. **a,** Bar plots showing the proportion of each cell type in young (Y), middle-aged (M), and old (O) ovaries estimated from snRNA-seq data. (Mean±SEM; Permutation test; *Padj<0.05, **Padj<0.01). **b,** Box plots showing the coefficient variation (CV) of ovarian cells at each age group. Box shows the median and the quartile range (25%–75%) and the length of whiskers represents 1.5× the IQR. (Wilcoxon test, **Padj < 0.01, ***Padj < 0.001). **c,** Box plots showing aging-associated transcriptional noise examined by CV analysis (O versus Y) in each cell type. Right shows the zoom-in view of the left panel. **d,** Upset plots showing the numbers of unique and shared pairwise differentially expressed gene (DEGs) between reproductive young mice and mice of other ages. **e,** Circos plots depicting the overlaps among gene lists of ovarian upregulated DEGs (left) or downregulated DEGs (right) for each cell type between old and young ovary. The inner-circle represents gene lists, and purple curves link identical genes. The genes that hit multiple lists are colored in dark orange, and genes unique to a list are shown in light orange. **f,** Box plots comparing the cosine similarities of transcriptomic changes during ageing between different age groups and tissues with human ovary (Wilcoxon test; *P<0.05, ***P<0.001, ****P<0.0001).

The change in the proportion of stromal cells and endothelial cells were similar, showing comparable abundance in middle age compared with the young ovaries but a significant decrease in the old ovaries **(Fig. 2a)**. Oppositely, the abundance of epithelial cells continued to increase during aging **(Fig. 2a)**, aligning with reported increases in thickness and outgrowth of ovarian surface epithelium in old ovaries^22^. Meanwhile, the proportion of immune cells in the ovary also underwent a significant temporal increase during aging **(Fig. 2a),** suggesting dysfunctional immune responses in the ovary throughout the aging process.

### Global transcriptional change during mouse ovarian aging

Given that increased transcriptional noise and perturbance are accompanied with aging^23,24^, we assessed transcriptional noise during ovarian aging by analyzing the expression differences of highly variable genes among cells of the same type within each age group and comparing the distribution of coefficient of variation (CV) between different age groups. Our findings revealed a significant increase in transcriptional noise in middle-aged and old ovaries compared to the young ovaries **(Fig. 2b and Fig. S1d)**. Subsequently, we evaluated aging-associated transcriptional perturbance by analyzing the expression changes of highly variable genes between individual young cells and individual old cells of the same type and then calculating the cell type-specific coefficient of variation. We discovered that granulosa cells and theca cells showed higher CV compared to other somatic cell types **(Fig. 2c)**. Similar results were found when comparing middle-aged ovary to young ovary **(Fig. S1e),** suggesting a higher vulnerability of cells involved in follicle development to reproductive aging than other somatic cell types.

To determine the change in gene expression during mouse ovarian aging, we first analyzed pairwise differentially expressed genes (DEGs) between reproductive young mice and mice of other ages. A total of 783 DEGs were identified when comparing ovaries between peri-estropause and reproductive young mice (M vs Y), with 317 upregulated and 466 downregulated **(Fig. 2d)**. A much more dramatic change occurred as mice approaching post-estropause (O vs Y), revealing 3668 DEGs, of which 1394 were upregulated and 2274 were downregulated. These results revealed a prevalence of downregulated DEGs during mouse ovarian aging, with a substantial increase in the number of DEGs when approaching post-estropause, suggesting a profound impact of estropause on the ovarian cells. Remarkably, 60% (192/317) of the upregulated DEGs and 77% (359/466) of the downregulated DEGs in peri-estropause ovaries remained detectable in post-estropause ovaries when compared to reproductive young ovaries. This finding suggested that most age-related gene expression changes observed in peri-estropause persist and contribute to the functional decay of the ovary in later life. Subsequently, we calculated the number of pairwise DEGs in each cell type **(Fig. S1f)** and found that LC, EC, and O had more DEGs in the peri-estropause stage compared to other cell types, whereas GC-A, LC, and SC exhibited more DEGs when approaching post-estropause.

Interestingly, numerous aging-associated DEGs, especially in the comparison between O vs Y, were shared among cell types **(Fig. 2e and Fig. S2a-b)**. To further explore the coordination of transcriptomic changes during ovarian aging, we calculated the pairwise cosine similarity of aging-associated DEGs in the ovary and compared it with the cosine similarity calculated from aging-associated DEGs in other tissues from the Tabula Muris Senis dataset. It was found that the mouse ovarian cells showed a significant increase in the coordination of transcriptomic changes during aging **(Fig. 2f)**. Moreover, mouse ovarian cells demonstrated a higher coordination value (Old vs Young) compared to other mouse tissues, even though the old ovary samples (15.5 months) were chronologically younger than those of other mouse tissues examined. **(Fig. 2f and Fig. S2c-k)**. This finding is consistent with our observations in human ovarian aging and further reinforces the notion that a high level of coordination is a distinctive characteristic of ovarian aging^25^.

### Monotonic transcriptomic signatures of mouse ovarian aging

To identify DEGs that constantly increased or decreased during ovarian aging, we employed loess regression to calculate the average trajectory for each gene within the union set of DEGs across different age groups and categorized these genes based on their expression patterns. Notably, due to the limited presence of only 8 oocytes in post-estropause samples, oocytes were excluded from the monotonic DEGs (MDEGs) analysis. Through this approach, we identified a total of 773 upregulated and 1679 downregulated DEGs exhibiting monotonic change during aging across 10 ovarian somatic cell types **(Fig. 3a)**.

**Figure 3.**
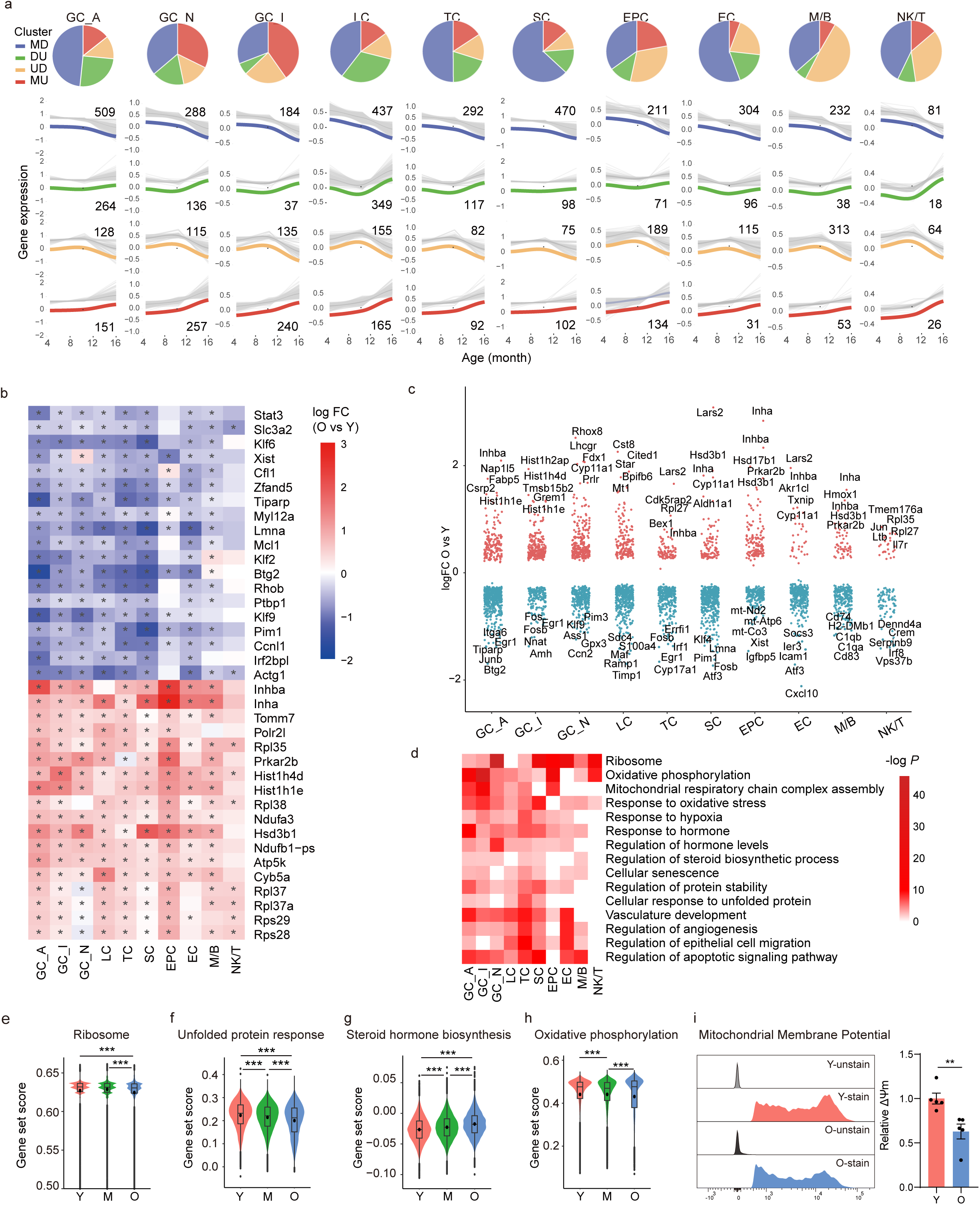
Monotonic temporal gene signatures of mouse ovarian aging. **a,** Pie chart showing the proportion of temporal DEGs with four different expression patterns during mouse ovarian aging (MD, monotonic down; DU, down and up; UD, up and down; MU, monotonic up) in each cell type (up panel). Gene trajectories during mouse ovarian aging and the number of genes within each expression pattern in each cell type (down panel). Gene expression was z scored, and trajectories of all genes were estimated by LOESS. **b,** Heatmap showing the log2 fold changes in gene expression (O vs Y) of monotonic upregulated DEGs shared by at least 6 cell types and monotonic downregulated DEGs shared by at least 7 cell types. * Indicates a statistically significant difference (Padj <0.05). **c,** Strip plots showing logFC (O vs Y) of monotonic DEGs in each cell type. Top five upregulated and top five downregulated genes per cluster labeled. **d,** heatmap showing the representative GO terms of monotonic DEGs in each cell type. **e-h,** Violin plots and box plots showing the gene set score of ribosome **(e)**, unfolded protein response **(f)**, steroid hormone biosynthesis **(g)** and oxidative phosphorylation **(h)** during mouse ovarian aging. Dot mark indicating the mean ***Padj < 0.001. **i,** Flow cytometry analysis of mitochondrial membrane potential in mouse ovarian cells. The Left Panel showed the representative histograms of mitochondrial membrane potential captured in the PE channel. The right panel showed the statistical comparison of average mitochondrial membrane potential between young and old groups. Mitochondrial membrane potential strengths were normalized to the young group. **, p<0.01; n=5.

Similar to the findings from pairwise DEG analysis, numerous MDEGs overlapped across different cell types **(Fig. S3a-b)**. Among them, 56 upregulated MDEGs and 189 downregulated MDEGs were commonly observed in at least 4 cell types. The most frequently upregulated genes included *Inhba*, *Inha*, and *Hsd3b1*, which are related to the biosynthesis of hormones such as inhibin, activin, and progesterone **(Fig. 3b)**. In addition, genes associated with ATP biosynthetic processes such as *Atp5k*, *Ndufb1*, *Atp5md*, *Ndufa* were also commonly upregulated. Moreover, we observed that *Lmna*, the gene encoding Lamin A/C, was consistently downregulated in more than 9 cell types. Lamins serve as structural components of the nuclear lamina, and genetic variants in *LMNA* have been reported to be associated with polycystic ovary syndrome and ovarian failure^26,27^. *Xist*, a non-coding RNA crucial for X chromosome inactivation, was also among the top shared downregulated MDEGs. Interestingly, the expression of *Xist* was reported to be downregulated with aging and upregulated following melatonin treatment in mouse ovary^28^ and was highly expressed in fetal mouse ovaries but sharply decreased after birth, contributing to perinatal oocyte loss through autophagy^29^. Other top frequently downregulated genes included *Stat3*, *Rhob*, *Klf6*, *Ccnl1*, and *Actg1*, mainly associated with developmental processes such as angiogenesis and response to growth factors, as indicated by GO enrichment analysis.

To delve deeper into cell type-specific DEGs undergoing monotonic changes, we calculated the log2 fold change (O vs Y) of each MDEG in each cell type. The top 5 upregulated and downregulated MDEGs in each cell type were highlighted in **Fig. 3c**. We observed that multiple genes belonging to the activator protein 1 (AP-1) superfamily of transcription factors, including *Fos*, *Fosb*, *Aft3*, and *Junb*, were consistently identified as the top downregulated monotonic DEGs across various cell types. Specifically, FOSB was recognized as a hub gene responsible for regulating the downregulated subset of genes in granulosa cells during primate ovarian aging^24^. Moreover, *Fos* null mice exhibited a failure to ovulate and form a corpus luteum (CL) even when administered exogenous gonadotropins, suggesting that ovarian expression of *Fos* is crucial for successful ovulation and CL formation^30^. The continuous downregulation of *Fosb* and *Fos* during ovarian aging implies that these genes might serve as key mediators in regulating ovarian function.

Notably, we found that certain cell type-specific downregulated MDEGs were marker genes or potential marker genes which were highly expressed in respective cell types, suggesting the loss of cell identity during ovarian aging. For example, *Cyp17a1* and *CD74*, the marker genes of theca cell and macrophage, respectively, were identified as top-downregulated MDEGs in corresponding cell types. To further investigate whether ovarian cells lose their cell identity during aging, we calculated the cell identity score in each cell type based on the expression of the top 50 cell type-specific genes. We observed that, during aging, most cell types exhibited a gradual decrease in the expression of their respective cell type-specific genes, except for GC-I and GC-N **(Fig. S3c).** These results suggest a prevalent loss of cell identity during mouse ovarian aging, which is similar to the findings in human ovarian aging^25^.

The Gene Ontology (GO) enrichment analysis of all MDEGs indicated common associations with pathways across different cell types (**Fig. 3d**). These pathways included oxidative phosphorylation, responses to oxidative stress, responses to hormones, regulation of protein stability, ribosome-related processes, and regulation of angiogenesis et.al. To further elucidate the effect of ovarian aging on these pathways, the pathway activity scores were examined. Results showed that the score of steroid hormone biosynthesis exhibited a continuous increase, whereas the score of ribosome-related signaling pathway and unfolded protein response consistently decreased during ovarian aging across almost all cell types (**Fig. 3e-g, Fig. S4a-c)**. Remarkably, the overall score of the oxidative phosphorylation (OXPHOS) pathway for all ovarian cells was significantly decreased during aging (**Fig. 3h**). However, the trends varied among different cell types, with an increase observed in GC-I, GC-N, and LC, while a decrease was noted in other cell types **(Fig. S4d)**. Dysfunction of mitochondrial OXPHOS has been reported in different organs during aging^31^. Interestingly, we observed a decrease in the gene expression of all mitochondrial-encoded genes, except for two bicistronic genes mt-*Nd4l* and mt-*Atp8***(Fig. S4e-f)**, alongside an increase in the gene set score of nuclear-encoded genes across most cell types during mouse ovarian aging **(Fig. S4g-h),** suggesting an imbalance between nuclear- and mitochondrially encoded OXPHOS subunits. To validate the OXPHOS function, we measured mitochondrial membrane potential and found a significant decline in aged ovarian cells **(Fig. 3i)**. These results indicated a disrupted mitochondrial homeostasis during mouse ovarian aging, with the downregulation of OXPHOS potentially attributed to the specific deficiency of mitochondria-encoded transcripts.

### Increased cellular senescence during mouse ovarian aging

Cellular senescence, an important hallmark of aging, has been found to increase in different tissues during aging^32^. To investigate if cellular senescence increased during mouse ovarian aging, we examined the cells expressing *Cdkn1a* (p21) in our single-cell data. The expression level of *Cdkn1a* was increased during aging in ovarian cells (**Fig. 4a**) and the most significant increase was found in GC-I (1.2-fold) and GC-N (1.8-fold) in old ovary compared with young ovary (**Fig. 4b**). Surprisingly, *Cdkn1a* expression was significantly decreased in LC from both middle-aged and old ovaries when comparing to young ovaries, while an increase of *Cdkn1b* (p27) expression level was observed during aging (**Fig. S5a**), suggesting that the cell cycle arrest of LC may be dependent on p27 instead of p21. To gain insight into the transcriptional signatures of *Cdkn1a*^+^ ovarian cells, we identified the DEGs between *Cdkn1a*^+^ cells and *Cdkn1a*^-^ cells. GO analysis revealed enrichment of these DEGs in signaling pathway such as cellular response to hypoxia, NF-kappa B signaling pathways, TNF signaling pathway, and regulation of protein stability **(Fig. S5b)**. Notably, key genes involved in NF-kB pathway such as *Nfkbiz*, *Nfkb1* and *Rela* were upregulated in *Cdkn1a*^+^ ovarian cells **(Fig. S5c)**. *Cdkn1a*^+^ cells exhibited a significantly higher NF-kB transcriptional activity compared to *Cdkn1a*^-^ cells across all cell types, as indicated by NF-kB pathway score (**Fig. 4c**). This suggests that NF-kB signaling-mediated inflammatory activation might be involved in cellular senescence in the mouse ovary.

**Figure 4.**
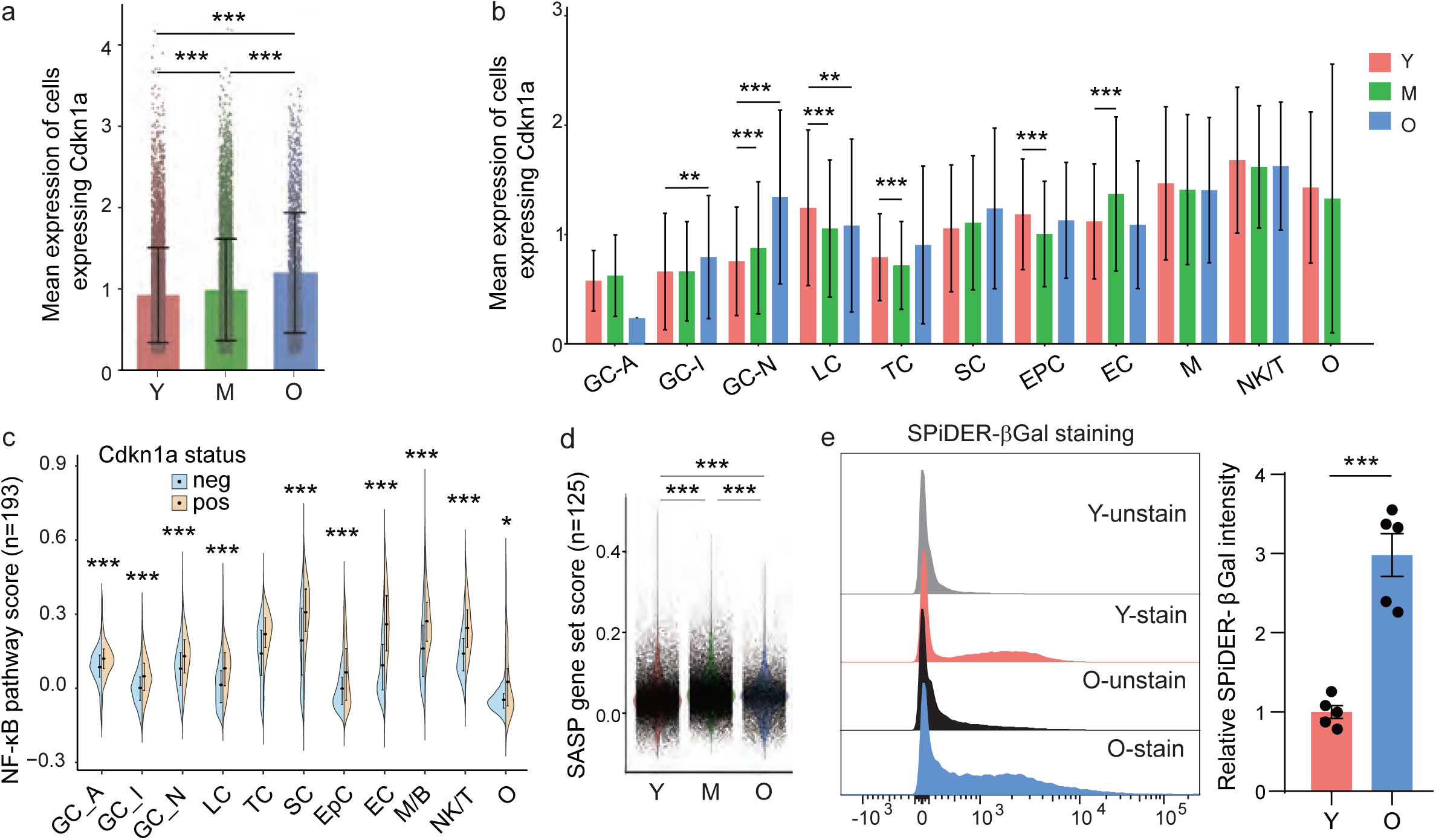
Increase of cellular senescence during mouse ovarian aging. **a,** Bar plots showing the expression of *Cdkn1a* for the cells that express the gene at each age group (Mean±SEM, Wilcoxon test, *** Padj < 0.001). **b,** Bar plots showing the expression of *Cdkn1a* in each cell type for the cells that express the gene at each age group. (Mean±SEM, Wilcoxon test, *** Padj < 0.001). **c,** Violin plots showing gene set score of NK-κB pathway in *Cdkn1a*+ cells and *Cdkn1a*-cells in each cell type. **d,** Violin plots showing the senescence-associated secretory phenotype (SASP) gene set score at each age group (Wilcoxon test, *** Padj < 0.001). **e** Flow cytometry analysis of SPiDER-β-Gal stained mouse ovarian cells. The Left Panel showed the representative histograms of the SPiDER-β-Gal signal captured in the GFP channel. The right panel showed the statistical comparison of SPiDER-β-Gal intensity between young and old groups. SPiDER-β-Gal intensity for each sample was calculated by subtracting the mean GFP intensity in the unstained sample from the mean GFP intensity in the stained sample. Background-removed intensities were normalized to the young group. ***, p<0.001; n=5.

Given that the senescence-associated secretory phenotype (SASP) in senescent cells is known to elicit chronic inflammation and contribute to organ aging, we next compared SASP gene set scores of all ovarian cells of different ages^33^. Significant increases in SASP gene set scores were observed in both middle-aged and old ovaries compared to young ovaries **(Fig. 4d),** while ovaries from peri-estropausal (M) showed the highest score. We subsequently examined the alterations in SASP scores across various cell types. Notably, GC-I and GC-N exhibited a consistent increase in scores throughout the aging process **(Fig. S5d)**. We further identified potential ovarian SASP marker genes by overlapping SASP-related genes and the monotonic upregulated DEGs during ovarian aging. Notable examples include *Igfbp4*, *Hmgb1*, and *Il-18*, which were recognized as canonical SASP genes, and consistently exhibited upregulation throughout the process of ovarian aging **(Fig. S5e)**. To validate the increase of cellular senescence burden in aged mouse ovaries, we performed SPiDER-βGal staining and noted increased signals of senescence-associated β-galactosidase (SA-β-gal) in live ovarian cells isolated from aged ovaries compared to that from young ovaries (**Fig. 4e**).

### Altered cellular communications during mouse ovarian aging

Aging is coupled with progressive alterations in intercellular communication^32^. To investigate the change of cell-cell communication throughout the reproductive lifespan of mice, we utilized comparative CellChat^34^ to analyze signaling interactions among all ovarian cell types at different ages. We found that the total number of possible interactions and the overall interaction strength progressively decreased during ovarian aging **(Fig. 5a-b and Fig. S6a)**. In the reproductively young ovary, GC-N, SC, and TC were the primary sources of incoming interactions, with SC and GC-N being the major contributors to outgoing interactions **(Fig. S6b)**. Both incoming and outgoing signals from these cell types decreased since peri-estropausal stage, eventually nearly disappearing, or dramatically reducing in the post-estropause ovary, especially for SC. These results indicate the importance of cell-cell interaction among granulosa cells, theca cells, and stromal cells in maintaining ovarian function. Conversely, the number and strength of interactions originating from and targeting epithelial cells and immune cells increased with ovarian aging (**Fig. 5b)**.

**Figure 5.**
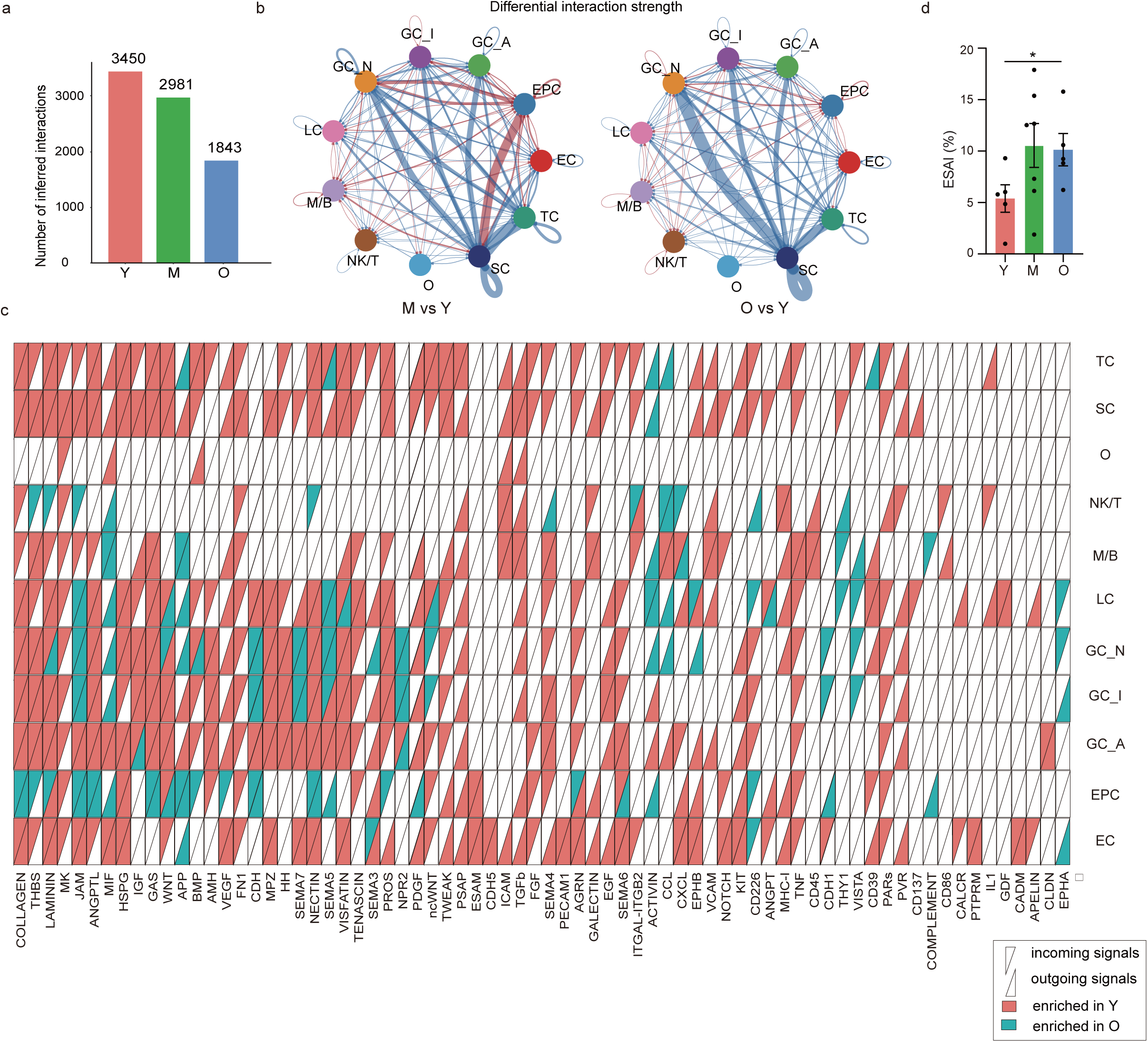
Decrease of cellular communications during mouse ovarian aging. **a,** Bar plots showing the number of intercellular interactions among ovarian cells at each age group. **b,** Circle plots showing the change of interaction strength in comparison between M versus Y (left panel) and between O versus Y (right panel). Blue lines indicate that the displayed communication is decreased in middle-aged (left panel) or old ovary (right panel), while red lines indicate that communication is increased in middle-aged (left panel) or old ovary (right panel). The arrows indicate the direction of intercellular communication. **c,** Heat map showing the outgoing and incoming signaling pathways significantly enriched in young or aged ovaries for each cell type. **d.** sEV secretion activity (ESAI) of each ovary sample from different groups. (Mean±SEM; *, p < 0.05).

To elucidate altered pathways of cell-cell communication during mouse ovarian aging, we conducted a comparative analysis of communication probabilities among all pairs of cell groups in the inferred networks of old ovaries and middle-aged ovaries compared to young ovaries **(Fig. 5c)**. We identified 73 and 72 pathways that exhibit a significant difference in communication probability in middle-aged ovary and old ovary when compared with young ovary, respectively **(Fig. S6c)**. Several pathways crucial for follicle development and maturation were significantly changed during aging. For example, the abundance of BMP signaling (Bmp15-(Bmpr1b+Bmpr2)), coming from oocytes and targeting granulosa cells, was the highest in young ovaries, decreased in middle-aged ovaries, and became entirely depleted in old ovaries **(Fig. S6d-e)**. The abundance of AMH signaling, primarily emanating from Amh-high granulosa cells, steadily declined with aging, aligning with the concurrent reduction in small follicles during the aging process **(Fig. S6f)**. Interestingly, we found that NPR2 signaling, which plays an essential role in maintaining oocyte meiotic arrest, was more abundant in aged ovaries. Further investigation into the expression patterns of the ligand (*Nppc*) and receptor (*Npr2*) across different age groups revealed an upward trend in *Nppc* expression and a decline in *Npr2* expression during aging. This dysregulation of NPR2 signaling may lead to abnormalities in oocyte meiosis, potentially contributing to the increased aneuploidy observed with reproductive aging **(Fig. S6g)**. Moreover, we found the MK signaling pathway, derived mostly from the stromal cells, significantly decreased during aging **(Fig. S6h)**. Mdk expression was mainly found in the stroma around the growing follicle but decreased around the antral follicle or corpus luteum^35^. Additionally, the supplement of midkine to early secondary follicles in vitro was shown to promote follicle growth^35^. These results indicate that the crosstalk between oocytes, granulosa cells, and stromal cells gradually lost during ovarian aging may contribute to the dysfunction of folliculogenesis and declined quality of oocytes.

Notably, the Collagen signaling and Thrombospondin (THBS) signaling, the core components of extracellular matrix (ECM) biology, showed significantly higher communication probability in most cell types in the young ovary, indicating the ECM disruption during aging **(Fig. 5c and Fig. S6i)**. However, epithelial cells exhibited a significantly higher communication probability of these signaling in the aged ovary. Collagen and THBS signaling, documented for their involvement in epithelial cell proliferation and migration^36,37^, could partially explain the observed increased proportion of epithelial cells and the reported outgrowth of ovarian surface epithelium in aging ovaries. Furthermore, signaling pathways related to inflammation and immune regulation, such as COMPLEMENT, VISTA (V-domain immunoglobulin suppressor of T cell activation), CCL, CD226, and THY1, were notably upregulated in aging ovaries, particularly in aged immune cells **(Fig. 5c)**. In summary, our results showed remarkable changes of cell-cell communication among different cell types during mouse ovarian aging, which may potentially contribute to dysfunction of follicle maturation, ECM disruption, inflammaging, and epithelial hyperplasia.

Apart from the intercellular communication mediated by direct ligand-receptor interaction, we evaluated indirect communication through small extracellular vesicle (sEV) secretion in mouse ovary during reproductive aging using SEVtras (sEV-containing droplet identification in scRNA-seq data), an algorithm capable of delineating sEV signals at droplet resolution and assessing sEV secretion activity across different cell types ^38^. SEVtras specifically detects signals from sEVs while remaining resilient to interference from cell debris and large EVs. We observed a significant increase in the sEV secretion activity index (ESAI) during reproductive aging **(Fig. 5d)**, consistent with the current knowledge that aging is accompanied by increased EV release partially attributed to elevated SASP, compromised immune response and inflammation and tissue damage^39^. After deconvoluting sEV secretion activity across various cell types based on the transcriptional similarity of origin cells with sEV and the sEV biogenesis capacity of origin cells, we found that EC, LC, IC, and TC exhibited stronger sEV secretion activity compared to other cell types **(Fig. S6j)**. Strikingly, EC contributed the most to the increased sEV secretion during reproductive aging. The function of EC-derived sEVs is not fully understood. However, given that EC serves as the interface between circulating blood and the vascular wall, they may communicate with cells from both sides, potentially impacting vascular integrity and inflammation.

### Changes in granulosa cells during mice ovarian aging

GC was the cell cluster with the most dramatic perturbations in transcriptome during mouse ovarian aging. Owing to their high cellular heterogeneity, we further subclustered GCs to examine the transcriptional changes in different GC types throughout the aging process. Unsupervised clustering divided GCs into 5 main subpopulations: pre-antral GCs (*Amh*^+^, *Gata*^+^), atretic GCs (*Itih5*^+^, *Cald1*^+^), mitotic GCs (*Top2a*^+^, *Mki67*^+^), cumulus GCs (*Ldha*^high^, *Cox4i2*^+^), mural GCs (*Nppc*^high^, *Mro*^+^) **(Fig. 6a-b)** ^40–42^. Trajectory analysis using Monocle 3 showed that mural GCs appeared the latest along the pseudotime when the pre-antral GCs were set as the root (starting point). This observation aligns with the stages of follicle development, as mural GCs are present during the formation of the antral follicle **(Fig. 6c-d)**. From pre-antral GCs to mural GCs, *Amh* expression gradually decreased and was almost undetectable in mural GCs, while expression of *Nppc*, *Cyp11a1*, *Cyp19a1*, and *Hsd3b1* steadily increased. Expression of *Fshr* and *Npr2* along with proliferation-related markers were highest at the middle stage where cumulus GCs and mitotic GCs were present. These changes in gene expression over the pseudotime corresponded with shifts in related hormone levels during follicle development.

**Figure 6.**
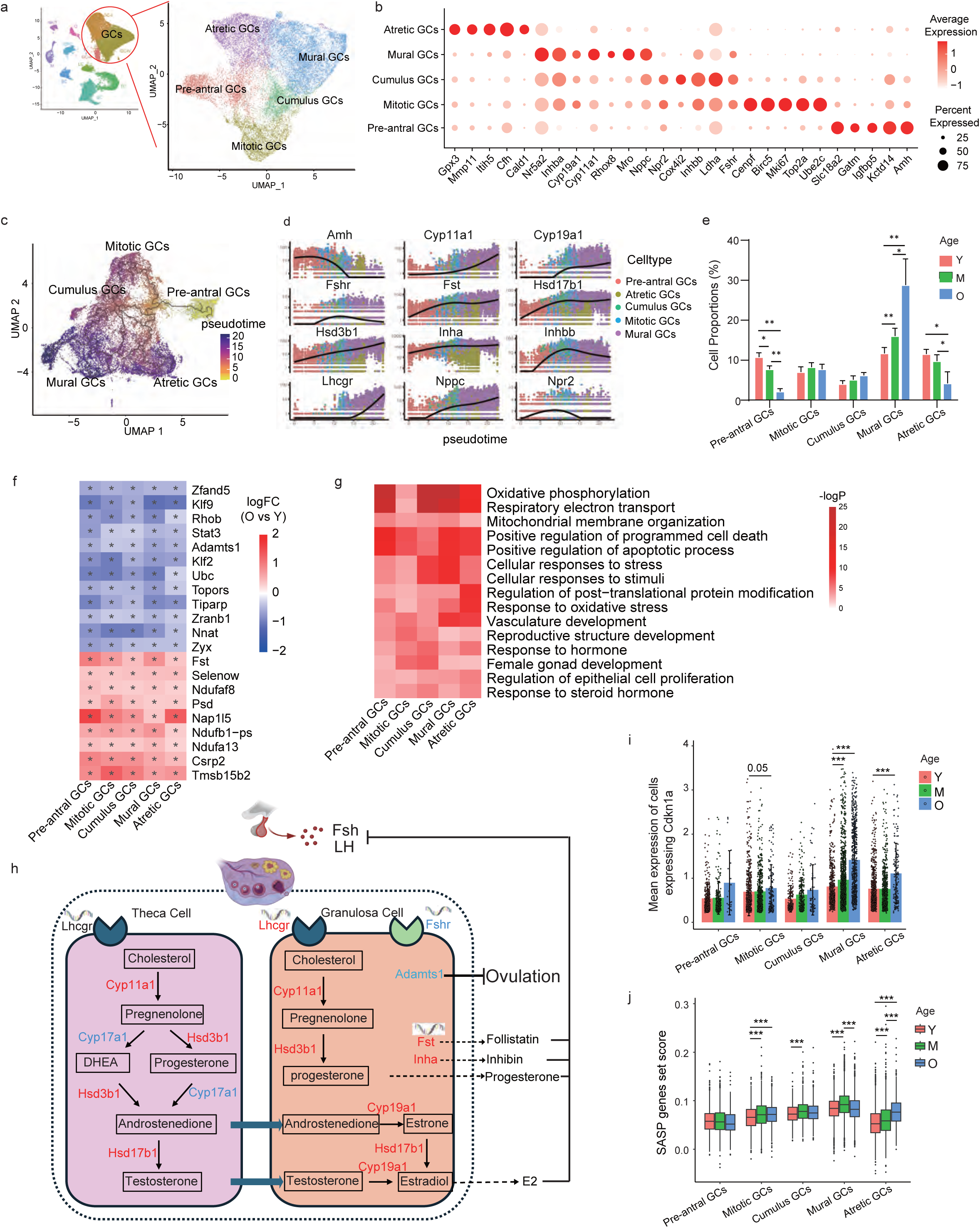
Change of subpopulations of granulosa cells during mouse ovarian aging. **a,** UMAP plots showing the subpopulations of mouse granulosa cells. **b,** Dot plots showing the expression of representative genes for each subpopulations of granulosa cells. **c,** UMAP plots showing pseudotime values of subpopulations of granulosa cells. **d,** Expression of hormone related genes in subpopulations of granulosa cells across pseudotime. **e,** Bar plots showing the proportion of each granulosa cell subpopulation in young, middle-aged, and old ovaries. (Mean±SEM; Permutation test; *Padj <0.05, **Padj<0.01). **f,** Heatmap showing the log2 fold changes in gene expression (O vs Y) of monotonic DEGs shared by all subpopulations of GCs. * Indicates a statistically significant difference (Padj <0.05). **g,** Heatmap showing the representative GO terms of monotonic DEGs in each subpopulation of GCs. **h,** Diagram showing ovarian steroidogenesis by granulosa cell and theca cell. Color of genes indicates upregulation (red) or downregulation (blue) of the gene expression in old ovary comparing to young ovary. **i,** Bar plots showing the expression of Cdkn1a in each subpopulation of GCs for the cells that express the gene at each age group. (Mean±SEM, Wilcoxon test, *** Padj < 0.001). **j,** Box plots showing the SASP gene set score in each subclusters of GCs at each age group.

Reproductive aging led to changes in the proportion of GC subpopulations. pre-antral GCs and atretic GCs were significantly decreased, while mural GCs were significantly increased during aging **(Fig. 6e)**, likely due to the exhaustion of early follicles in the old ovaries. Interestingly, we found that the increase in the proportion of mural GCs during aging was largely attributed to the increase of *Lhcgr*^+^ mural GCs in the post-estropause ovary **(Fig. S7a-b)**. Consistently, the expression level of *Lhcgr* in mural GCs dramatically increased in the old ovary compared with young or middle-aged ovary, which was one of the top 3 upregulated DEGs of old mural GCs along with *Rhox8* and *Cyp11a1* (**Fig. S7c-e)**. Typically, mural granulosa cells express high levels of *Lhcgr* exclusively in preovulatory follicles, in contrast to theca cells which constitutively express *Lhcgr*^43^. In PCOS ovaries, higher expression of LH receptor is observed in granulosa cells at early stages of follicular development, leading to premature luteinization of granulosa cells^44^. Additionally, studies have found upregulated *LHCGR* and *CYP11A1* in granulosa cells from older infertile patients (ages 43-47) from IVF when compared with young (ages 21-29) and middle-aged oocyte donors (ages 30-37), suggesting the age-related decline in female fertility is associated with the premature luteinization of granulosa cells occurring in older women^45^. Further investigation is needed to determine whether the increase in *Lhcgr* expression or *Lhcgr*+ cells contributes to the age-related decline in mouse fertility.

To further unravel the transcriptional change in GC subpopulations during aging, we identified the MDEGs within each subcluster **(Fig. 6f)**. We found a consistent decrease in *Adamts1*, a gene crucial for normal ovulation, across multiple GC subpopulations. Impaired ovulation due to mature oocytes remaining trapped in ovarian follicles has been documented in *Adamts1*-null mice^46^. Additionally, we found increased expression of *Fst,* which encodes follistatin, a glycoprotein that binds activin and decreases FSH secretion. Elevated follistatin levels have been associated with an increased risk for type 2 diabetes and polycystic ovary syndrome (PCOS) in human^47^. GO analysis of all MDEG in granulosa cells during ovarian aging revealed common associations with pathways involved in oxidative phosphorylation, cellular response to stress, vasculature development, regulation of epithelial cell proliferation, and response to hormone **(Fig. 6g)**. As key producers of ovarian steroids such as estradiol and progesterone, granulosa cells, in cooperation with theca cells, regulate the production of FSH and LH from the pituitary gland through both negative and positive feedback mechanisms. We further investigated the change of hormone synthesis-related genes in GCs during aging and found common changes across different subpopulations of GCs, including the decrease of *Fshr*, and the increase of *Cyp11a1*, *Hsd3b1*, *Cyp19a1*, *Hsd17b1*, and Inha **(Fig. 6h and Fig. S7f)**. Additionally, dysregulation of hormone-related genes was also found in theca cells, including decreased levels of *Cyp17b1* and increased levels of *Hsd3b1*, *Hsd17b1*, and *Cyp11a1*. Notably, the observed trends of changes in hormone-related genes during mouse ovarian aging contrasted with those observed during human ovarian aging based on human ovarian single-nuclear RNA-seq data **(Fig. S7g)**. These differing observations between humans and mice may explain the differences in hormone fluctuations between post-estropausal mice and post-menopausal women. It has been reported that post-estropausal mice generally maintain moderate to high circulating levels of 17β-estradiol and progesterone but relatively lower levels of LH and FSH, whereas post-menopausal women exhibit very low levels of 17β-estradiol and progesterone but significantly elevated levels of FSH and LH^6^.

Moreover, we found that all GC subpopulations showed concordant increases in *Cdkn1a* expression **(Fig. 6i)**. Significant changes were noted in the mitotic, mural, and atretic GCs, especially in the post-estropause ovary. Furthermore, all GC subpopulations, except for pre-antral GC, showed a significant increase in SASP score **(Fig. 6j)**. These findings indicate that granulosa cells undergo senescence consistently during aging, which could play a vital role in contributing the ovarian aging.

### Changes in stromal/ Theca cells during mice ovarian aging

Ovarian stromal cells constitute a heterogenous cell population, serving a dual role by providing structural support for follicle development and engaging in intricate bidirectional paracrine signaling with the follicle^48^. Theca cells were previously believed to be recruited from surrounding stromal tissue during folliculogenesis^49^. However, the nature of cellular heterogeneity and the differentiation trajectory within theca cells (TCs) and stromal cells (SCs), as well as the specific impact of ovarian aging on SCs/TCs subpopulations, remains elusive.

Here, mouse SCs/TCs were classified into 8 subpopulations by unsupervised clustering, encompassing interstitial stromal cell (ISC, *Dcn*^+^, *Lum*^+^, *Kcnk2*^+^), smooth muscle cell (SMC, *Acta*^+^, *Actg*^+^), pericyte (P, *Rgs5*^+^, *Ebf1*^+^), 2 groups of early theca cells (*Enpep*^+^, *Hiip*^+^), and 3 groups of steroidogenic theca cells (*Star*^+^, *Aldh1a1*^+^) **(Fig. 7a-b)**. Transcriptional markers for mitotic early theca cells (MET) and non-mitotic early theca cells (NMET) were found to be similar, except for proliferation-related markers such as *Top2a*, *Mki67*, and *Ube2c*, which were specifically expressed in MET.

**Figure 7.**
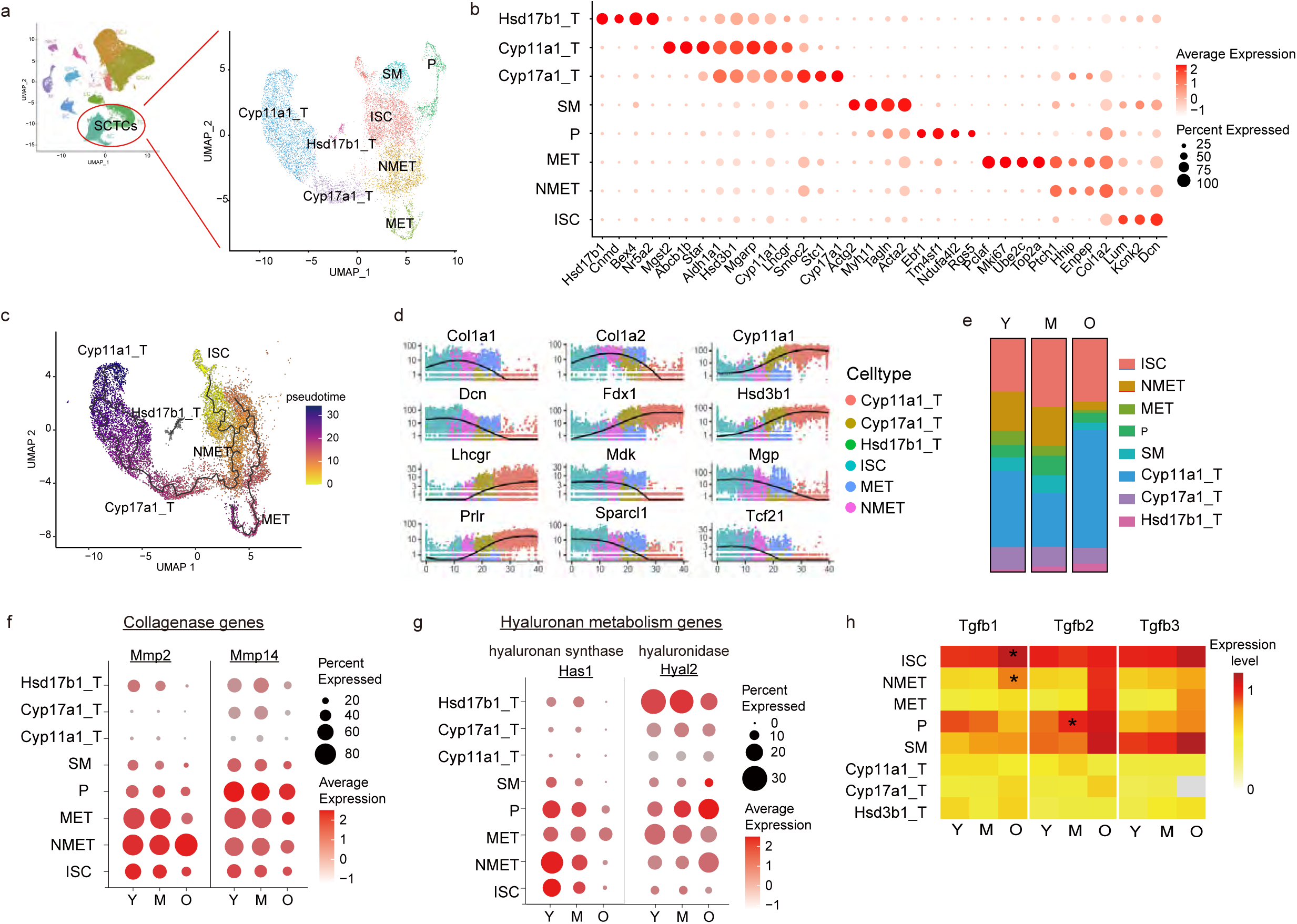
Change of subpopulations of stromal and theca cells during mouse ovarian aging. **a,** UMAP plots showing the subpopulations of mouse stromal and theca cells. **b,** Dot plots showing the expression of representative genes for each subcluster of stromal/theca cells. **c,** UMAP plots showing pseudotime values of subpopulations of granulosa cells. **d,** Kinetics plots showing the relative expression of representative genes for SC/TC subclusters along pseudotime. **e,** Bar plots showing the distribution of subpopulations of stromal/theca cells types at each age. **f,** Dot plots showing the expression of collagenase genes in subclusters of SC/TCs. **g,** Dot plots showing the expression hyaluronan metabolism genes in subclusters of SC/TCs. **h,** Heatmap showing the expression of Tgfb genes in each subclusters of SC/TCs at each age group. *Padj < 0.05 when comparing with young ovary.

These *Enpep*^+^ early theca cells were also enriched for *Ptch* and *Thbd*, which are marker genes identified in human perifollicular stromal cells^50^. These perifollicular stromal cells have been reported to be located directly adjacent to the GC basement membrane in growing follicles, which may play a pivotal role not only in regulating follicular development through secretory signals but also in their ability to differentiate and replenish theca cells. The steroidogenic theca cells can be further divided into 3 subgroups based on the representative steroidogenic enzyme markers: Cyp17a1-high theca cell, Cyp11a1-high theca cell, and Hsd17b1-high theca cell. *Cyp17a1*, *Cyp11a1*, and *Hsd17b1* are genes encoding the key enzyme catalyzing different processes in androgen synthesis in theca cells. These findings suggest the potential existence of subtypes of highly differentiated cell populations in the theca layer with specialized steroidogenic function, collectively contributing to androgen genesis.

To better elucidate the relationship between ISCs and theca cells and changes in gene expression over the theca cell differentiation trajectory, we performed trajectory analysis using Monocle 3 **(Fig. 7c-d)**. Based on the developmental pseudotime, NMETs could function as precursors of steroidogenic theca cells. NMETs may originate from ISCs and subsequently differentiate into Cyp17a1-high theca cells and Cyp11a1-high theca cells. METs could be a group of self-renewal early theca cells also originating from ISCs but without differentiation potential. From ISCs to *Cyp11a1*-high theca cells, the gene expression of extracellular matrix-associated genes (*Dcn*, *Mgp*, *Sparcl1*, *Tcf21*) gradually decreased while steroidogenic and steroid hormone receptor-related genes (*Cyp11a1*, *Hsd3b1*, prolactin receptor (*Prlr*), *Lhcgr*, *Fdx1*) gradually increased. Moreover, the expression of Mdk gradually decreased and was associated with the acquisition of steroidogenic TC cell fate, suggesting that ISCs and early theca cells play vital roles in the secretion of midkine to promote follicle development. The change of cell composition within the SCTC cluster was characterized by the exhaustion of early theca cells in the old ovary **(Fig. 7e)**. Early theca was thought to form the theca interna of secondary and preantral follicles while steroidogenic theca was thought to form the theca interna of antral follicles^19^. The marked decrease in the proportion of early theca cells during mouse ovarian aging is consistent with the depletion of growing follicles post-estropause.

Increased stiffness, an aging-related change in the old ovary, depends on stromal ECM including collagen and hyaluronan^51^. Excessive collagen accumulation was reported to be the primary cause of increased stiffness in many tissues including the ovary. We found stromal cell is the major cell type robustly expressing collagen-associated genes in mouse ovary **(Fig. S8a)**. Collagen genes expressing Type I, II, IV, V, and VI collagens are the most abundant **(Fig. S8b)**. We further looked at the expression of these relatively highly expressed collagen genes in stromal subtypes and found collagen genes were modestly expressed in steroidogenic theca cells, but elaborately expressed in interstitial stromal cells, early theca, and pericyte **(Fig. S8c)**. Collagen gene expressions in ISCs were unexpectedly decreased while they were well maintained in early theca and pericyte during reproductive aging. We also checked the gene expression of collagenases that break the collagen and found *Mmp2* was uniquely expressed in SCs and *Mmp14* was ubiquitously expressed in SCs and many other cell types **(Fig. S8d)**. We observed the loss of *Mmp2* and *Mmp14* expression in ISCs and MECs in aged mice **(Fig. 7f)**, suggesting the failure of break collagens may led to the accumulation of collagen in old ovaries. Since hyaluronan acid (HA) reduction is another mechanism causing increased stiffness, we next focused on HA metabolism genes including three hyaluronan synthases (*Has1*, *Has2*, and *Has3*) and four hyaluronidases (*Hyal1*, *Hyal2*, *Cemip*, and *Cemip2*). SCs showed a relatively high expression of *Has1* although Has genes were low abundant in all ovarian cell types **(Fig. S8e)**. Interestingly, we found hyaluronidase gene *Cemip* was dominantly expressed in luteal cells, which may explain why HA was minimal in the corpus luteum (CL)^51^. We further compared the changes of hyaluronan metabolism genes during reproductive aging and found the decline of *Has1* in ISCs, pericyte, and NMET as well as the increase of *Hyal2* in pericyte and NMET, suggesting the dysregulated HA synthesis and degradation in aged ovary **(Fig. 7g)**. Furthermore, significantly increased expression of *Tgfb1*, a central mediator of collagen deposition, hyaluronan synthesis, and fibrogenesis, in ISC and NMET in the old ovary was observed, suggesting their contribution to fibrogenesis during mouse ovarian aging **(Fig. 7h and Fig. S8f-h).**

### Changes in immune cells during mice ovarian aging

Since the immune cell clusters in mouse ovaries showed mixed cell type signatures, we further performed unsupervised clustering and identified a total of 19 clusters from the initial two immune cell clusters **(Fig. 8a-b)**. Each cluster exhibited a unique gene expression pattern indicative of a distinct cell type. In addition to lymphoid cells like B cells (*Cd79a*^+^), T cells (*Cd3*^+^), natural killer cells (NKs, *Klrb1c*^+^), and innate lymphoid cells (ILCs, *Il1rl1*^+^), we identified diverse myeloid cell populations with high-resolution, including macrophages (Mφs, *Cd14*^+^), type 1 conventional dendritic cells (cDC1s, *Xcr1*^+^ and *Clec9a*^+^), type 2 conventional dendritic cell (cDC2s, *Tnip3*^+^ and *Cd209a*^+^), plasmacytoid dendritic cells (pDCs, *Siglech*^+^), neutrophils (*S100a8*^+^ and *S100a9*^+^) and mast cells (*Cpa3*^+^) **(Fig. 8b)**. Among these cell populations, macrophages displayed the greatest diversity with 8 subtypes which will be further characterized. We also identify two distinct subsets of cDCs, cDC1s and cDC2s were markedly separated on UMAP. cDC2s, known to share the macrophage markers and to activate ILCs and T cells, is closer to Mφs on UMAP **(Fig. 8a)**. cDC1s, capable of cross-presenting cell-associated antigens, can be divided into two subsets **(Fig. S9a)**. One subset expressed classical cDC1 markers *Xcr1* and *Clec9a*, while the other expressed low *Xcr1* and *Clec9a* but high *Ccl22*, *Cacnb3*, *Nudt17*, *Mreg*, *Acdy6* and *Il4i1* **(Fig. S9a)**. A previous study showing similar cDC1 subsets in colon cancer has suggested that *Ccl22*-cDC1s may represent activated cDC1s rather than a developmentally distinct subset^52^. Our findings highlight the presence of diverse immune cell types within the mouse ovary.

**Figure 8.**
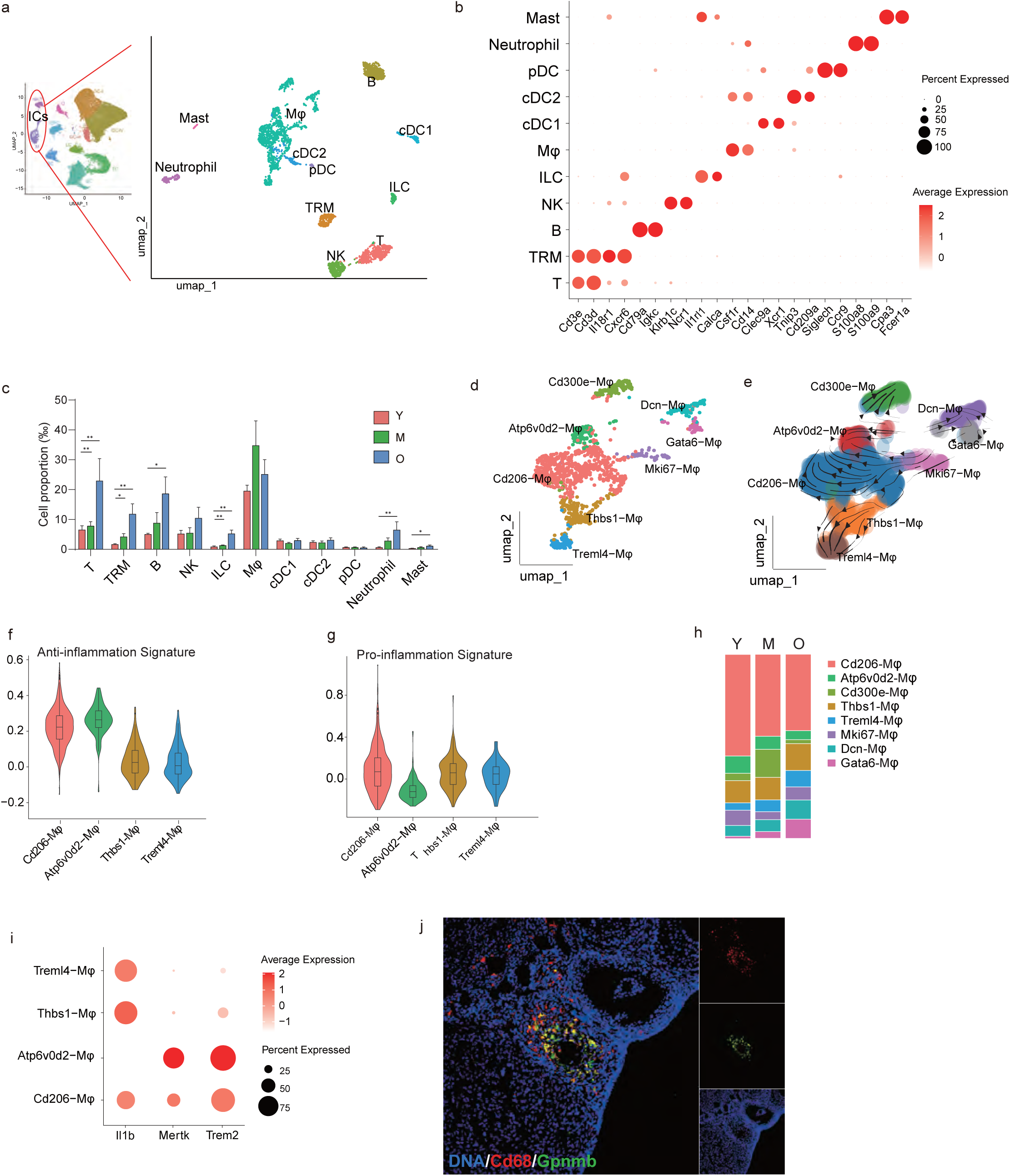
Change of subpopulations of immune cells during mouse ovarian aging. **a,** UMAP plots showing the subpopulations of mouse immune cells. **b,** Dot plots showing the expression of representative genes for each subcluster of immune cells. **c,** Bar plots showing the proportion of each immune cell types in young, middle-aged, and old ovaries. (Mean±SEM; Permutation test; * Padj<0.05, **P<0.01). **d**. UMAP plots showing the subpopulations of macrophages. **e.** RNA velocity stream for macrophages projected onto the UMAP embedding. **f**. Violin plots showing M2 anti-inflammatory gene set score in different macrophages. **g**. Violin plots showing M1 pro-inflammatory gene set score in different macrophages. **h**. Proportion of different subpopulations of macrophage among macrophage cluster. **i**. Dot plot showing the expression of inflammatory gene Il1b and endocytosis gene Mertk and Trem2 in subcluster of macrophages. **j**. Representative image showing Gpnmb-positive macrophages in young ovaries.

During reproductive aging, we observed a gradually accumulated proportion of T, B, NK, ILC, neutrophil and mast cell, while the proportion of total macrophages was exceptionally unchanged **(Fig. 8c)**. Our focus then turned to the eight macrophage clusters identified through unsupervised cluster analysis, namely Cd206*-*Mφ (*Mrc1*^+^, *Pf4*^+^), Atp6v0d2-Mφ (*Atp6v0d2*^+^, *Gpnmb*^+^), Cd300e-Mφ (*Cd300e*^+^, *Asb2*^+^, *Fcrl5*^+^), Thbs1-Mφ (*Thbs1*^+^, *Ccr2*^+^), Treml4-Mφ (*Treml4*^+^, *Ace*^+^), Mki67-Mφ (*Mki67*^+^, *Top2a*^+^), Dcn-Mφ (*Dcn*^+^, *Mgp*^+^), Gata6-Mφ (*Gata6*^+^, *Esr2*^+^) **(Fig. 8d and S9b)**. Utilizing RNA velocity analysis, which models mRNA transcription, splicing, and degradation kinetics^53^, we inferred cellular state transitions among these subpopulations. This analysis revealed a clear directional flow from proliferative Mki67-Mφ to Cd206-Mφ and then diverged into either Atp6v0d2-Mφ or Thbs1- and Treml4-Mφ **(Fig. 8e)**. Using signature genes of “Pro-inflammatory” M1 and “Anti-inflammatory” M2 macrophages^54^, we found the co-expression of both M1 and M2 gene signatures in Cd206-Mφ and Cd300e-Mφ **(Fig. 8f-g)**, consistent with previous studies^55^, suggesting a more complex phenotype of *in vivo* macrophage compared to *in vitro* activation state of macrophages. Notably, Treml4- and Thbs1-Mφ exhibited strong pro-inflammatory signatures, while Atp6v0d2-Mφ exhibited strong anti-inflammatory signatures **(Fig. 8f-g)**. We also noticed the proportion of pro-inflammatory Treml4- and Thbs1-Mφ increased with reproductive aging, while the proportion of anti-inflammatory Atp6v0d2-Mφ decreased **(Fig. 8h)**. To understand the potential function of Atp6v0d2-Mφ, we compared its gene expression with that of other macrophages and found genes highly expressed in Atp6v0d2-Mφ were enriched in functions including phagocytosis **(Fig. S9c)**, suggesting Atp6v0d2-Mφ was a type of macrophage with phagocytotic capacity. Consistently, the expression of phagocytosis-associated genes including *Mertk* and *Trem2* were stronger in Atp6v0d2-Mφ compared to other macrophages, while the pro-inflammatory gene *Il1b* was almost absent **(Fig. 8i and S9d)**. During ovarian aging, we saw a trend of declined expression of Mertk and Trem2 in Atp6v0d2-Mφ **(Fig. S9e)**, suggesting a loss of phagocytotic capacity in this macrophage. We further verified the presence of phagocytotic Mφ in the corpus luteum, as well as in potential atretic follicles or cysts, where apoptotic or defective cells are abundant **(Fig. 8j and S9f)**. Taken together, we identified a unique macrophage in mouse ovary involved in phagocytosis and both the cell proportion and phagocytosis capacity of this cell type declined during reproductive aging.

### Transcriptomic signatures of estropausal transition at periestropausal stage

To unravel the molecular and cellular mechanism that may drive the transition of cycle irregularity, we compared the ovaries from irregular cycling mice (M-ir) with regular cycling mice (M-r) during the estropausal transition stage. No significant change was observed in cell composition between M-ir and M-r ovaries **(Fig. 9a)**. However, the transcriptional perturbation was significantly higher in mice with irregular cycle comparing to mice with regular cycle, particularly in GC-A, TC, EPC, EC, and NK/T cell types **(Fig. 9b-c)**. Granulosa cells were shown to be the most vulnerable cell types affected by cycle irregularity, which is consistent with the impact of aging **(Fig. 9d)**.

**Figure 9.**
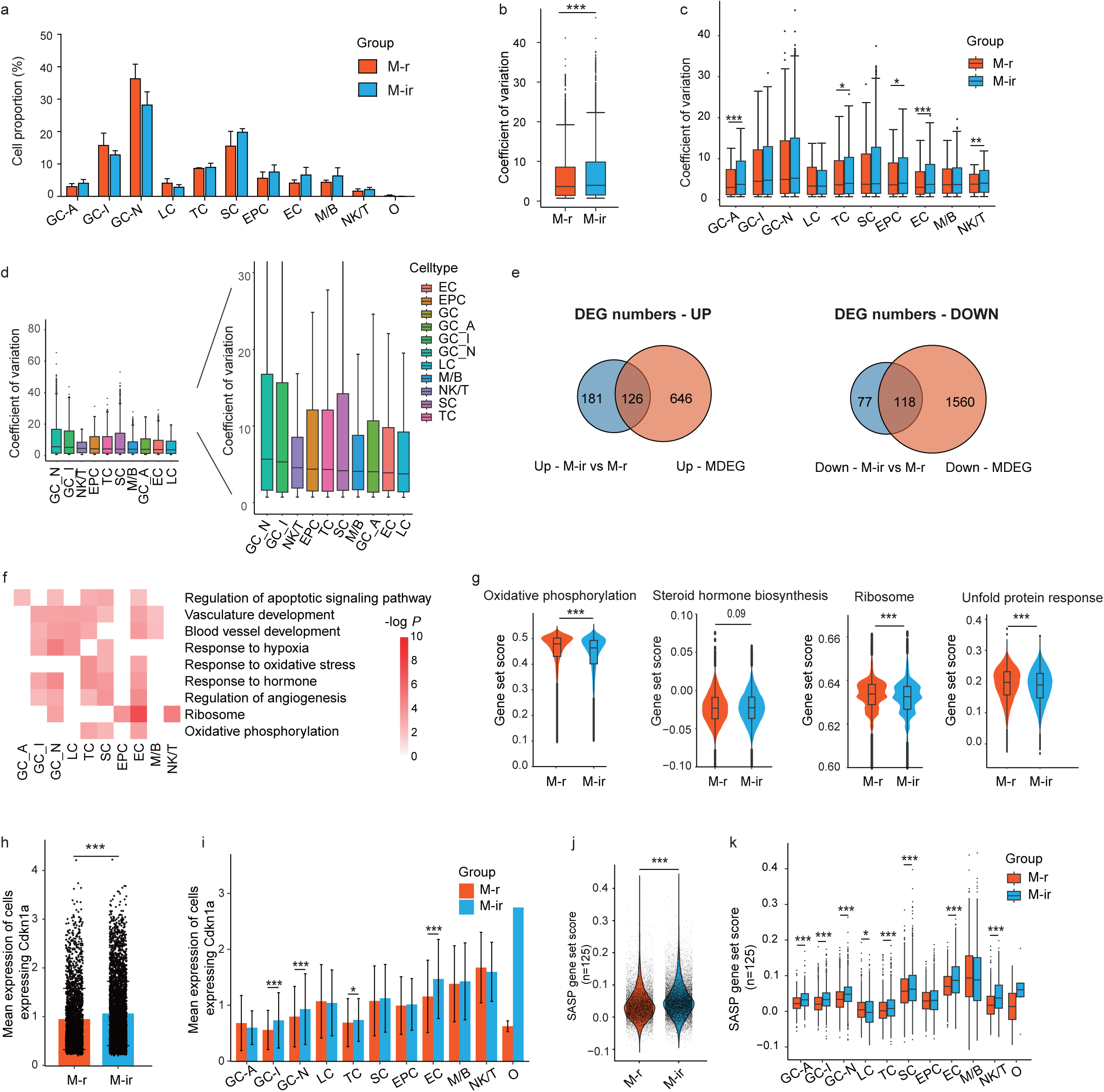
Transcriptome difference between mice with irregular cycle and regular cycle during estropausal transition stage. **a,** Bar plots showing the proportion of each cell type in middle-aged ovaries with regular cycle (M-ir) and irregular cycle (M-ir) (Mean±SEM). **b,** Box plots showing the coefficient variation (CV) of ovarian cells in irregular and regular cycling mice at transition age (Wilcoxon test, ***Padj < 0.001). **c,** Box plots showing the CV of each cell type in irregular regular cycling mice at transition age (Wilcoxon test, *Padj < 0.05, **Padj < 0.01, ***Padj < 0.001). **d,** Box plots showing cycle prolongation-associated transcriptional noise examined by CV analysis in each cell type. Right shows the zoom-in view of the left panel. **e,** Venn plots showing the overlaps of DEGs between the comparison of M-ir versus M-r and the monotonic DEGs observed during aging. left panel showing the upregulated DEGs, while right panel showing the downregulated DEGs. **f,** Heatmap showing the representative GO terms of DEGs between irregular cycling ovary and regular cycling ovary. **g,** Box plots and violin plots showing the gene set score of oxidative phosphorylation, steroid hormone biosynthesis, ribosome and unfolded protein response in irregular and regular cycling ovary at transition age (Wilcoxon test, ***Padj < 0.001). **h,** Bar plots showing the expression of Cdkn1a for the cells that express the gene in irregular and regular cycling ovary at transition age (Mean±SEM, Wilcoxon test, ***Padj < 0.001). **i,** Bar plots showing the expression of Cdkn1a for the cells that express the gene in each cell type in irregular and regular cycling ovary at transition age (Mean±SEM, Wilcoxon test, ***Padj < 0.001). **j,** Violin plots showing the SASP gene set score of ovarian cells at M-r and M-ir group. **k,** Violin plots showing the SASP gene set score of each cell type at M-r and M-ir group (Wilcoxon test, *Padj < 0.05, ***Padj < 0.001).

Next, we performed DEG analysis and identified 307 upregulated and 195 downregulated DEGs comparing irregular cycling mice with regular cycling mice. Among them, 41% of the upregulated DEGs and 61% of the downregulated DEGs overlapped with aging-related upregulated or downregulated MDEGs **(Fig. 9e)**. GO analysis indicated the enrichment of DEGs in pathways including response to oxidative stress, response to hormone, oxidative phosphorylation, ribosome and blood vessel development across different cell types **(Fig. 9f)**. We subsequently computed pathway scores and observed that the changes on pathways in mice with irregular cycles compared to those with regular cycles exhibited consistent trends with the changes observed during ovarian aging. This included a decrease in oxidative phosphorylation, ribosome, and unfolded protein response pathways, alongside an increase in steroid hormone biosynthesis in irregular cycling mice compared to regular cycling mice **(Fig. 9g)**. Additionally, the expression levels of *Cdkn1a* were significantly elevated in irregular cycling mice compared to regular cycling mice, particularly in GCs, TCs, and ECs **(Fig. 9h-i)**. Moreover, mice with irregular cycles also showed significantly higher SASP scores, specifically in GCs, SCs, TCs, ECs, and NK/T cells **(Fig. 9j-k)**. These results indicated that the ovarian aging-associated signatures in mice were recapitulated in mice with irregular cycles when comparing to mice with regular cycles at the same chronological age. Our findings suggest that aging-associated molecular changes in the ovary may drive the transition of cycle irregularity and lead to estropause in mice.

## Discussion

In the present study, we characterized aging signatures in mouse ovaries across the reproductive lifespan ranging from reproductively young (regular cycle) through peri-estropause (regular vs. irregular cycles) to post-estropause age (acyclic) at single-cell resolution. Additionally, we uncovered the molecular alterations in mouse ovaries associated with the onset of cycle irregularity during the peri-estropausal stages by comparing single-cell transcriptomic data between irregular and regular cycling ovaries in middle age. To our knowledge, our study is the first to provide an in-depth understanding of cell type-specific mechanisms of mouse ovarian aging across the reproductive lifespan concomitant with cycle changes at single-cell resolution.

The stages of reproductive senescence, from regular cycling to irregular cycling to acyclicity, are common features between the preclinical rodent model and women, which provides researchers with opportunities to gain a fundamental understanding of the key elements underlying reproductive aging. Our study characterized the signature of mouse ovarian aging along with age-related cycle pattern changes. We conducted daily monitoring of the estrous cycle in reproductive young, peri-estropausal, and post-estropausal mice, respectively, over a minimum period of 3 weeks. Surprisingly, we found that the cycle length of most reproductive young mice exceeded 4-5 days until they reached 4.5-month-old. This finding is consistent with previous reports that C57BL/6J mice showed a U-shaped distribution of cycle length variance with age, while the majority of mice started to show 4-5 days cycle length around 4-5 months^9^. During the estropausal transition stage, approximately 50% of mice showed a lengthening of cycling, indicative of irregular fertility and impending reproductive senescence. Around 15.5 months, some mice exhibited acyclicity, indicating entry into the post-estropausal stage. Aging-related changes in the mouse ovary, including increased transcriptomic heterogeneity, p21, and SASP gene expression and decreased cellular communications have already been significantly shown since middle-age. Upon estropause with acyclicity, a more substantial alteration in mouse ovarian transcriptomics was observed, concomitant with the collapse of ovarian function in aged mice. The trigger for the transition to menopause remains a topic of debate, with some theories suggesting it could be initiated by a decline in the follicle pool, dysregulation of the hypothalamic-pituitary-gonadal (HPG) axis, or a combination of both^6^. It has been reported that a complex set of changes in neuroendocrine and neurotransmitter signaling involving hypothalamic GnRH neurons and alterations in glutamatergic, GABAergic, and monoaminergic signaling, likely play roles in the early stages of the transition to a reproductively senescent state for rodents, non-human primates, and women alike^56,57^. We found that despite identical genetic background, environment, and chronological age, irregular cycling mice showed multiple aging-associated molecular changes in ovarian somatic cells comparing to regular cycling mice, including higher transcriptional heterogeneity, upregulation of senescence marker gene expression, and dysregulation of aging hallmarks pathways. Our findings suggest that the aging-related changes in ovarian somatic cells may also play a vital role in the onset of cycle irregularity and estropausal transition. Moreover, it has been reported that transplanting oocyte-depleted ovaries from young donors into reproductively aged mice (at 17 months of age) can extend lifespan and alleviate circulating inflammation in the recipients^58^. This underscores the pivotal role of ovarian somatic cells in preserving somatic health within the organism. In conjunction with our findings, this suggests that targeting pathways related to aging hallmarks in ovarian somatic cells may delay the onset of estropause (menopause) and confer broader health benefits.

Mitochondrial dysfunction is one of the hallmarks of aging and potentially drives ovarian aging^59^. Abnormalities in mitochondrial ultrastructure and integrity, metabolism, dynamics, and mtDNA mutations and deletions have been associated with aging of oocytes and granulosa cells^60,61^. Mitochondrial oxidative phosphorylation (OXPHOS) system, which uses enzymes to oxidize nutrients and thereby release energy for ATP production, is central to the cellular energy supply. Inefficiencies in the OXPHOS system may result in the production of high levels of ROS, leading to cellular dysfunction and apoptosis^62^. The nuclear and mitochondrial genomes each encode different subunits of the electron transfer chain of the OXPHOS system and closely coordinate their activities. Our study revealed that overall OXPHOS was downregulated in the aged ovaries. Further investigation indicates that the decline in OXPHOS observed in aged ovaries is primarily due to the loss of mitochondria-encoded electron transport chain (ETC) components but not nuclear-encoded components. Similar to our findings, it was reported that during mouse skeletal muscle aging, there was a specific loss of mitochondrial, but not nuclear, encoded OXPHOS subunits^63^. Additionally, they found that knockout of SIRT1 mimics aging by decreasing mitochondrial, but not nuclear-encoded, OXPHOS components. Taken together, the imbalance between nuclear- and mitochondrially-encoded OXPHOS subunits could be a potential mechanism underlying the loss of membrane potential and mitochondrial homeostasis during ovarian aging.

Senescent cell burden has been shown to increase with age in various tissues during aging and aging-related disease. We identified an elevated senescence burden in ovarian cells during reproductive aging, characterized by upregulated expression of *Cdkn1a* and SASP genes. This was further confirmed through flow cytometry, demonstrating an increased signal of SA-β-gal. Overexpression of SA-β-gal in senescent cells serves as a widely recognized non-genetic marker of cellular senescence. In our attempt to quantify β-gal in mouse ovaries using SPiDER-βGal staining, we encountered difficulties in identifying SA-β-gal positive cells via fluorescence microscopy due to strong autofluorescence in aged ovaries, which is attributable to the lipofuscin accumulation. The lipofuscin-like autofluorescence (lipo-AF) exhibited a dramatic increase during reproductive aging in the ovary, particularly in macrophage multinucleated giant cells (MNGC)^7^. A similar accumulation of lipo-AF has been observed in brain aging, specifically in microglia, the resident macrophage of the central nervous system (CNS)^64^. Apart from the MNGC with large nuclei located in follicle-free regions showing strong lipo-AF, we also found the diffuse distribution of lipo-AF-positive cells with small nuclei within ovaries. Consistently, flow cytometry analysis of unstained single ovarian cells excluding large cells like MNGC demonstrated that autofluorescence cells increase dramatically during reproductive aging. lipo-AF undermined the reliability of fluorescence-based assays for assessing aged ovaries. To address this limitation, we compared background-removed SPiDER-βGal intensities rather than the percentage of SA-β-gal-positive cells between cells isolated from young and old ovaries and demonstrated the increased SA-β-gal activity in non-MNGC somatic ovarian cells in aged ovaries. It would be interesting to investigate if these lipo-AF-positive cells are more prone to being senescent.

Our study highlighted the cell type-specific changes during mouse ovarian aging, particularly in granulosa cells, stomal/theca cells, and immune cells. Granulosa cells were the most affected somatic cell types during mouse ovarian aging. Tarin et al. proposed that a reduced ability of oocytes and granulosa cells to counteract reactive oxygen species (ROS) was among the most important physiological inducers of cellular injury associated with ovarian aging^14^. All sub-populations of granulosa cells in our data showed enrichment of aging pathways such as oxidative phosphorylation and response to oxidative stress. Consistently, previous studies reported the increase of oxidatively damaged lipids, proteins, DNA, and oxidative stress signaling in older granulosa cells ^65^. Moreover, our results showed that almost all kinds of granulosa cells showed increase of cellular senescence markers including *Cdkn1a* (p21) and SASP genes. In line with these results, it was reported that ROS induced an increase of p16, p21, and SA-β-gal activity in granulosa cells in vitro^66^. These findings suggest that dysregulated oxidative phosphorylation in aged ovaries may lead to the increase of ROS, which in turn induces cellular senescence in aged granulosa cells.

The theca that surrounds growing follicles, along with stromal cells, are critical modulators of the extracellular matrices remodeling process during ovulation and aging. Folliculogenesis and ovulation undergo repeated cycles of connective tissue remodeling and wound healing^22^. The continuous remodeling that takes place since puberty is thought to be one of the causes of fibrosis in aged ovaries due to increased collagen synthesis and decreased deposition of hyaluronan in both mouse and human samples^22,51^. However, recent studies present conflicting findings. Ouni et al. conducted a detailed analysis of mechanical matrisome in human ovarian biopsies across different ages up to menopause and reported no age-related differences in collagen and glycosaminoglycans, while the relative amount of thick collagen fibers diminished^67^. Meanwhile, Winkler et al did Picro-Sirius red staining on all female reproductive tissues in mice at different ages and observed collagen accumulation in all female reproductive tissues except for the ovary and cervix^18^. In our single-cell transcriptomic analysis, we found no changes in collagen gene expression during aging but noted decreases in collagenase genes, specifically *Mmp2* and *Mmp14*, in ISCs and mitotic early theca cells. Consistently, a previous single-cell study also reported no change in collagen expression and a decrease in *Mmp2* expression in fibroblasts comparing 9-month-old ovaries with 3-month-old ovaries^17^. However, contrary to previous findings that reported significant upregulation of the fibrogenesis regulators *Tgfb1* and *Tgfb2* in theca cells of 9-month-old ovaries by IPA analysis^17^, our data showed significant upregulation of *Tgfb1* expression in ISC and NMET, and significant upregulation of *Tgfb2* expression in pericytes. Furthermore, our data revealed a significant decrease in the expression of the hyaluronan synthase gene *Has1* in NMET, P, and ISC, while the hyaluronidase gene *Hyal2* showed a predominant increase in P, potentially contributing to the decline in hyaluronan levels observed in the aged ovary. Taken together, our results suggest that ISC, early theca cells, and pericytes may play crucial roles in ovarian fibrogenesis during aging, though the exact mechanisms warrant further investigation.

Increased immune cells, especially T cells, is a common age-related change in different aged tissues including ovaries^17,18,68^. In line with other studies, we also observed increased lymphoid cells in ovaries during reproductive aging. Moreover, we focused on the myeloid cells. We identified macrophages, cDC1s, cDC1s, pDCs, and mast cells in mouse ovaries even though the cell number was low, suggesting the ovary contains diverse myeloid cells presented in most other tissues. Moreover, we focused on the biggest population which was macrophage among myeloid cells. Interestingly, we found changes in macrophage subtypes during reproductive aging though the proportion of macrophages showed no significant change. Based on RNA velocity analysis, we identified the development path from immature macrophages to either pro-inflammatory macrophages or phagocytotic macrophages. Decreased phagocytotic macrophages and increased pro-inflammatory macrophages were observed during reproductive aging. Notably, a unique macrophage subtype showing almost absent expression of the pro-inflammatory gene *Il1b* and strong expression of phagocytosis gene *Mertk* was identified. This macrophage highly expressed *Atp6v0d2* and *Gpnmb*. *Atp6v0d2* is a macrophage-specific V-ATPase subunit in the lysosome, and it restricts inflammasome activation and bacterial infection by facilitating lysosome fusion with autophagosome or phagosome^69^. In addition, Atp6v0d2-macrophages are recruited to injured nerves and are required for efficient removal of myelin through phagocytosis in a sciatic nerve injury model, suggesting the important role of Atp6v0d2-macrophages in tissue damage repair^70^. *Gpnmb* is also a phagocytic protein that controls the trafficking of cellular apoptotic debris for degradation and is essential for kidney repair^71^. More importantly, decreased *Mertk* expression in this macrophage suggested the decline of phagocytic capacity and impaired tissue repair during reproductive aging. Whether enhancing the clearance capacity of this macrophage subtype could potentially counteract ovarian aging requires further exploration. Moreover, MNGCs are believed to result from the fusion of macrophages with accumulating debris in aged tissues^7^. Since Atp6v0d2-macrophage is the only subtype with phagocytosis transcriptional marks, it would be interesting to study if MNGC comes from these Atp6v0d2-macrophages.

In summary, our study provided an in-depth understanding of global and cell type-specific mechanisms underlying mouse ovarian aging across the reproductive lifespan concomitant with cycle changes at single-cell resolution. We observed significant remodeling of the ovarian cellular architecture with aging, marked by a decline in Amh-high granulosa cells, theca cells, stromal cells, and endothelial cells, coupled with an increase in epithelial cells and immune cells. Additionally, aging was associated with increased global transcriptional noise and decreased cell identity in ovarian cells. Genes undergoing monotonic changes across cell types during reproductive aging were consistently enriched in pathways related to aging pathways such as oxidative phosphorylation, stress responses, and proteostasis. Furthermore, senescence marker gene expression, including *Cdkn1a* (p21), and senescence-associated secretory phenotype (SASP) factors were heightened during ovarian aging, particularly in granulosa cells. Aging also led to a significant decrease in cell-cell communications, particularly between stromal and granulosa cells, and an increase in extracellular vesicle secretion in mouse ovaries. Moreover, we identified age-related cell type-specific changes, including dysregulation of hormone synthesis in granulosa cells, alteration of collagen and hyaluronan metabolism in stromal and early theca cells, and functional decline of a unique phagocytosis-associated tissue-resident macrophage. Importantly, most of these aging-associated molecular changes were more pronounced in irregular cycling ovaries at the estropausal transition age compared to regular cycling counterparts, suggesting that aging-related molecular alterations in the ovary drive cycle irregularity and estropausal transition in mice. Thus, our findings provide insights into reproductive aging-associated biomarkers and potential cellular and molecular targets for improving aging-associated ovarian disorders, serving as a currently lacking resource for the field.

## Materials and Methods

### Mouse colony

The 3-month-old, 9.5-month-old, and 14.5-month-old female C57BL/6 mice were all virgin and obtained from the Jackson laboratory. All mice were housed in the SPF environment and in groups of up to 5 mice per cage with a 12-hour light /12-hour dark cycle. Mice had ad libitum access to water and food. All colonies were regularly controlled for infections using sentinel mice to ensure a healthy status. All experiments were approved by the ethical committee of Columbia University.

### Estrous cycle staging cytology

Vaginal smears were collected every day before noon using a pipette containing PBS and inserted in the vagina of the restrained mouse. Mucous tissue was then trickled on glass slides and visualized under the microscope to determine the estrous cycle. The cellular composition of the smears was analyzed according to known cell distribution patterns. The regular estrous cycle was defined as every 4 to 5 days and consists of four phases, which was proestrus, estrus, metestrus, and diestrus. The irregular cycle was defined as 2 contiguous cycles of 5–8 days. Acyclicity was defined as no cycle lasting more than 9 days.

### Cell suspension and library preparation for Single-cell RNA-seq

Ovary samples were collected separately from each individual at different days. Estrous cycle stages of mice were measured before noon and mice at diestrus stage were anesthetized and perfused with PBS. Two ovaries were isolated and immersed in chilled M2 medium (Sigma, M7167). Surrounding fat and connective tissue were trimmed and removed. Two intact ovaries were minced in 1.5 ml Eppendorf tube and treated by enzymatic digestion in dissociation buffer (M2 medium supplemented with 30 ug/ml Liberase DH (Roche, LIBDH-RO) and 1000U/mL DNase (Thermo Fisher)) at 37°C for 15 min with shaking on thermomixer (1000 rpm). The tube was briefly spun at 1000 rpm and the supernatant was collected in a 15 ml centrifuge tube. Next, the remaining tissues were cut and incubated in a dissociation buffer for 10 min with shaking. The cell suspension was pooled with the supernatant in a 15 mL tube and neutralized with M2 medium with 1% BSA. The pooled cell suspension was filtered with a 70-um cell strainer (Miltenyi Biotec) and was centrifuged. Cell pellets were incubated with RBC Lysis Buffer (Thermo Fisher) to remove red blood cells and subsequently washed with 1%BSA in PBS and 10% FBS in EMEM. Pelleted cells were resuspended in 10% FBS in EMEM and further submitted to Columbia Genomic Core for scRNA-seq library construction. Briefly, the cell suspension was loaded onto Chromium Single Cell Chip from 10x Genomics Chromium Single Cell 3’ Reagent Kits v3, aiming for a target output of 5000 cells per sample. Followed by reverse transcription, cDNA amplification, indexed adaptor ligation, and library amplification were performed according to the manufacturer’s protocol. The final libraries were sequenced on Illumina NovaSeq 6000 at Columbia Genome Center.

### Processing and quality control of scRNA-seq data

Raw FASTQ reads were processed using Cellranger analysis pipeline (version 5.0.1) (10× Genomics). Cellranger count command was used to align reads to the mouse genome reference (mm10 - 2020-A) and generate the gene expression matrices. The filtered gene-barcode count matrices were used for quality control with R package Seurat (version 4.1.0)^1^. Specifically, quality control filtered out cells with fewer than 200 genes, cells with more than 6000 genes, or cells with more than 15% mitochondrial genes by Seurat “*subset*” function. Additionally, we identified potential doublets introduced by technical artifact in each sample using DoubletFinder (version 2.0.3)^2^ and excluded the doublets by Seurat “*subset*” function. After integration and clustering, one ambient RNA cluster lacking specific gene markers and with low gene content was discarded.

### scRNA-seq data analysis and cell-type identification

The R package Seurat (version 4.1.0)^1^ was used for scRNA-seq data analysis, including data normalization, dimensionality reduction, cell clustering, and differential expression gene (DEG) analysis. Gene counts were normalized with the LogNormalize method using the Seurat function *“NormalizeData”* and the top 10% highly variable genes (HVGs) identified by the Seurat function *“FindVariableFeatures”* were used for principal component analysis (PCA) using the Seurat function *“RunPCA”*. To remove batch effects, the first 15 PCs were batch corrected using Harmony (version 0.1)^3^. Clustering was performed by constructing a K-nearest neighbor (KNN) graph with corrected PCs and applying the Louvain algorithm. Dimensional reduction was performed with Uniform ManifoldApproximation and Projection (UMAP) and individual clusters were annotated based on expression of cell type-specific markers. Cell type-specific markers were identified with the Seurat function *“FindAllMarkers”* for genes detected in at least 25% of cells and a log fold-change (LogFC) threshold of 0.25 using the non-parametric two-sided Wilcoxon rank-sum test.

### DEG

Cell-type specific pairwise differential expression analysis was performed with the Seurat function “*FindMarkers*” to identy DEGs between Y and M, between Y and O, and between M_ir and M_r in each cell type. The log fold change and adjusted P-value of each DEG were calculated by the MAST test and only those expressed in at least 25% cells with |“avg_logFC”| > 0.25 and “pval_adj” < 0.05 were considered to be DEGs. Gene ontology analysis was performed using Metascape. Scores of cell identity, pathways, and gene sets were evaluated by the Seurat function *“AddModuleScore”* with corresponding gene lists (Supplemental Table 1), respectively.

Cell-type specific dynamic differential expression analysis was performed by following steps: 1) Cell-type specific pairwise DEGs between Y and O, Y and M as well as M and O were merged to get a list of whole DEGs; 2) gene expression of these whole DEGs were z scored and locally estimated scatterplot smoothing (LOESS) regression was fitted for each gene to obtain predicted age-based gene expression values; 3) genes were further categorized into four clusters as monotonic-up (MU), up-down(UD), down-up (DU) and monotonic-down(MD) according to their changes from Y to M to O. MU and MD genes were regarded as genes undergoing monotonic changes during reproductive aging.

### Coefficient of variation analysis between different groups

Analysis of age-relevant CV was used to observe the aging effects on different cell types as described previously^4,5^. The Seurat “*FindVariableFeatures”* function was used to identify highly variable genes (HVGs). The top 10% variable genes (2744 of 27,445 genes) were selected for downstream analysis. Next, the absolute value of the cell-paired-distance dc,x was calculated for each HVG expression between all cells from two different groups in each cell type c.

To compare between young group (cell number = y) and the reproductively aged group (cell number = O), the formula is:

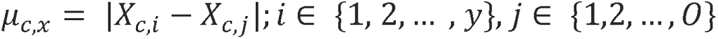

To compare between young group (cell number = y) and the reproductive transition group (cell number = k), the formula is:

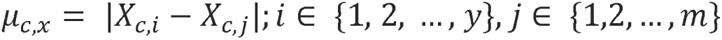

To compare between regular cycling group (cell number = r) and irregular cycling group (cell number = ir) at the same reproductive transition age, the formula is:

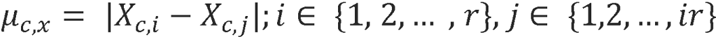

Next, the arithmetic mean of dc,x (mc,x), and the standard deviation of dc,x as (sc,x) were calculated. Accordingly, the transcriptional variation of each cell type between two different groups is defined by the following formula:

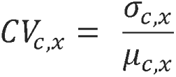

### Transcriptional heterogeneity analysis

The Seurat “*FindVariableFeatures”* function was used to identify highly variable genes (HVGs). The top 10% variable genes (2744 of 27,445 genes) were selected for downstream analysis. The absolute value of the cell-paired-distance dc,x was calculated for each HVG expression within each cell type c (cell number = n), the formula is:

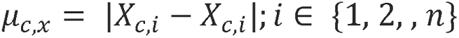

Next, the arithmetic mean of dc,x (mc,x), and the standard deviation of dc,x as (sc,x) were calculated. Accordingly, the transcriptional heterogeneity of each cell type in a specific group is defined by the following formula:

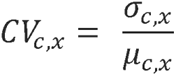

### Pairwise cosine similarity

For mouse ovary, the middle aged and reproductively old ovary were compared to the reproductively young ovary to identify the aging associated DEGs. For other mouse tissues, the data were obtained from Tabula Muris Senis^6^, using the 3-month group as a young reference, and compared it with both early-aged (18-month) and late-aged (24- or 30 month) groups. For human ovary, data were obtained from Chen et al.’s study^7^. The analytical approach of DEGs in mouse ovary were utilized to data of other mouse tissues and human ovary. The log2(fold changes) of aging associated DEGs in each cell type were extracted. Pairwise cosine similarity between all cell types within each tissue was computed using the “cosine” function in lsa package (v0.73.2). This generated a similarity matrix per tissue. The mean cosine similarity for other mouse tissue was statistically compared to the mean similarity in mouse ovary cell types using a two-sided Wilcoxon rank sum test.

### Gene set variation analysis (GSVA)

R package GSVA (v1.44.5)^8^ was utilized to estimate pathway activity scores for each cell, based on KEGG pathways in the MSigDB database (R package msigdbr v7.5.1). The GSVA score was normalized with the z-score method and used to indicate the pathway activity.

### Trajectory Analysis

The R package Monocle 3 (version 1.3.1)^9^ package was applied to construct pseudotime trajectories and to identify genes that play key roles during cell state transition in granulosa cells, thecal, and stromal cells. The original embedding of UMAP reduction was used. Monocle algorithm learns the sequence of gene expression changes each cell must go through as part of a dynamic biological process and places each cell at its proper position in the trajectory. The construction of single-cell trajectories was performed using default parameters. After learning the trajectory graph, Amh-high granulosa cells in the GC subset and interstitial stromal cells in the SCTC subset were chosen as the root node to order cells along the trajectory. Significantly changed genes along the pseudotime trajectory were identified using the Moncle3 graph-autocorrelation analysis to find genes that vary over the trajectory.

### RNA velocity analysis

To perform the RNA velocity analysis, we rerun the Cellranger count command with the argument --include-introns true. The aligned bam files possorted_genome_bam.bam generated by Cellranger were used to count the spliced reads and unspliced reads by the velocity (version 0.17.17)^10^. The RNA velocity estimation and visualization as streamlines were done by using the scvelo (version 0.3.1).

### SEVtras analysis

The secretion of small extracellular vesicles was analyzed using SEVtras (version 0.2.8, small extracellular vesicles(sEV)-containing droplet identification in scRNA-seq data)^11^. The raw_feature_bc_matrix folder generated by the Cellranger count command was used as input in *SEVtras.sEV_recognizer* function with default parameters to identify sEV-containing droplets. With the output of SEVtras.sEV_recognizer, sample- or cell type-specific sEV secretion activity indexes (ESAI) were calculated using *SEVtras.ESAI_calculator* function with default parameters.

### Cell-cell communication

Cell-cell communication analysis was performed using the R package CellChat (Version 1.5.0)^12^. based on the expression of known ligand-receptor pairs in different cell types. Mouse ligand-receptor interaction database (CellChatDB.mouse) were used to identify over-expressed ligands or receptors in each cell type. Cell-cell communication network was inferred based on the over-expressed ligands-receptors interactions if either ligand or receptor are over-expressed. Significant cell-cell communication was inferred by assigning each interaction with a probability value and peforming a permutation test.

### SPiDER-**β**-Gal staining

Cellular Senescence Detection Kit (SG04, Dojindo) was used to detect SA-β-Gal activity according to the manufacturer’s manual. For analysis by flow cytometry, mouse ovaries were processed using the same method for single-cell RNA-seq to obtain single-cell suspension. Cells were incubated with Bafilomycin A1 working solution in an M2 medium at 37 °C for 1 hour to inhibit endogenous β-galactosidase activity. The SPiDER-β-Gal working solution was further added to stain cells at 37 °C for 30 minutes. Cells were washed with M2 medium and analyzed by BD LSR flow cytometer. We calculated the background-removed SPiDER-β-Gal intensity for each sample by subtracting the mean GFP intensity in the unstained sample from the mean GFP intensity in the stained sample. Subsequently, these background-removed intensities were normalized to the young group.

### Mitochondrial membrane potential measurement

Mitochondrial Membrane Potential Detection Kit (MT13-10, Dojindo) was used to detect Mitochondrial membrane potential in ovarian cells according to the manufacturer’s manual. Mouse ovaries were processed using the same method for single-cell RNA-seq to obtain single-cell suspension. Cells were incubated with MT-1 Dye in M2 medium at 37 °C for 30 minutes and washed with M2 medium. Cells were resuspended in 1x Imaging Buffer solution and analyzed by BD LSR flow cytometer. Unstained samples were utilized to establish the positive cutoff on the PE channel and the mean intensity of PE-positive cells in stained sample was calculated to represent the mitochondrial membrane potential of each sample. Subsequently, the intensities were normalized to the young group.

### Immunofluorescence staining

Ovaries were freshly collected from mice at diestrus, fixed in 4% paraformaldehyde, embedded in Tissue-Tek O.C.T. Compound (Sakura Finetek). and sectioned at 10 μm thickness. Tissue sections were permeabilized in 0.04% Triton X-100/PBS for 15 mins. After blocking with 10% donkey serum (Jackson ImmunoResearch Labs)/PBS for 1 h, sections were incubated with primary antibodies at 4 °C overnight and the corresponding secondary antibody (Invitrogen) at RT for 45 mins, slides were mounted using ProLong Gold Antifade Mountant (Thermo Fisher). Nuclei were stained with Hoechst 33342 (62249, Thermo Fisher). Primary antibodies were listed below: anti-Gpnmb (R&D, AF2330), and anti-Cd68 (Abcam, ab53444). Images were captured using the Leica Stellaris 8 Confocal Microscope.

