## Supplemental Figures and Figure Legends for "Transcriptomic signatures of mouse ovarian aging and estropausal transition at single cell resolution"

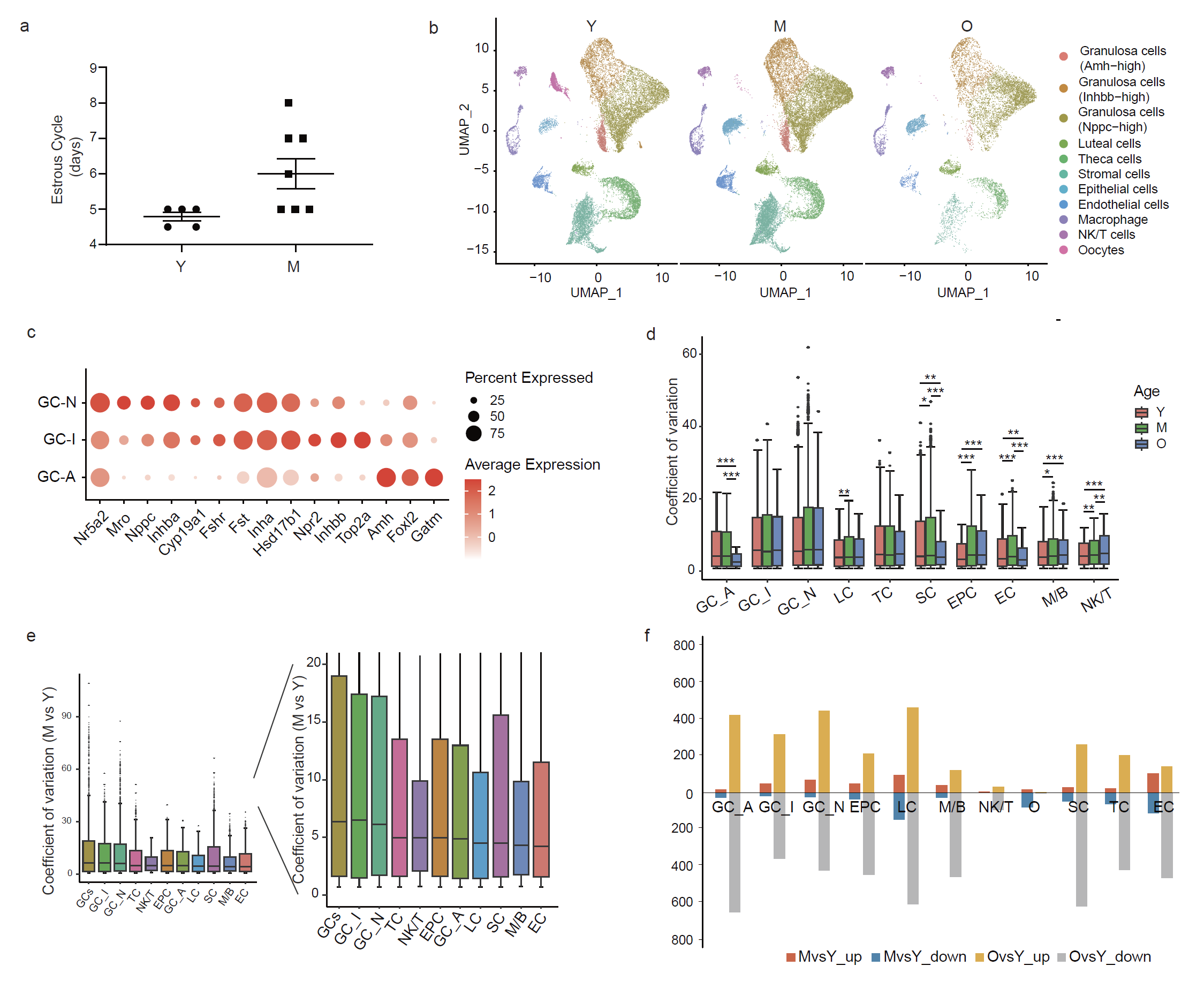

**Supplementary figure 1. Estrous cycle status of ovarian samples and global transcriptional change during ovarian aging. a,** Scatter plots showing the estrous cycle length of each young and middle-aged ovary submitted for single-cell sequencing. old ovaries showing acyclicity were submitted for scRNA-seq. **b,** UMAP plots showing mouse ovarian cells at each age. **c,** Dot plots showing the expression of representative genes for GCs. GC-A, Amh-high granulosa cell; GC-I, Inhbb-high granulosa cell; GC-N, Nppc-high granulosa cell; LC, luteal cell. **d,** Box plots showing the coefficient variation (CV) of each cell type at each age. Box shows the median and the quartile range (25%–75%) and the length of whiskers represents 1.5× the IQR. (Wilcoxon test, *Padj < 0.05, **Padj < 0.01, ***Padj < 0.001). **e,** Box plots showing aging-associated transcriptional noise examined by CV analysis (M versus Y) in each cell type. The right shows the zoom-in view of the left panel. **f,** Bar plots showing the numbers of pairwise differentially expressed genes (DEGs) in each cell type between reproductive young mice and mice of other ages.

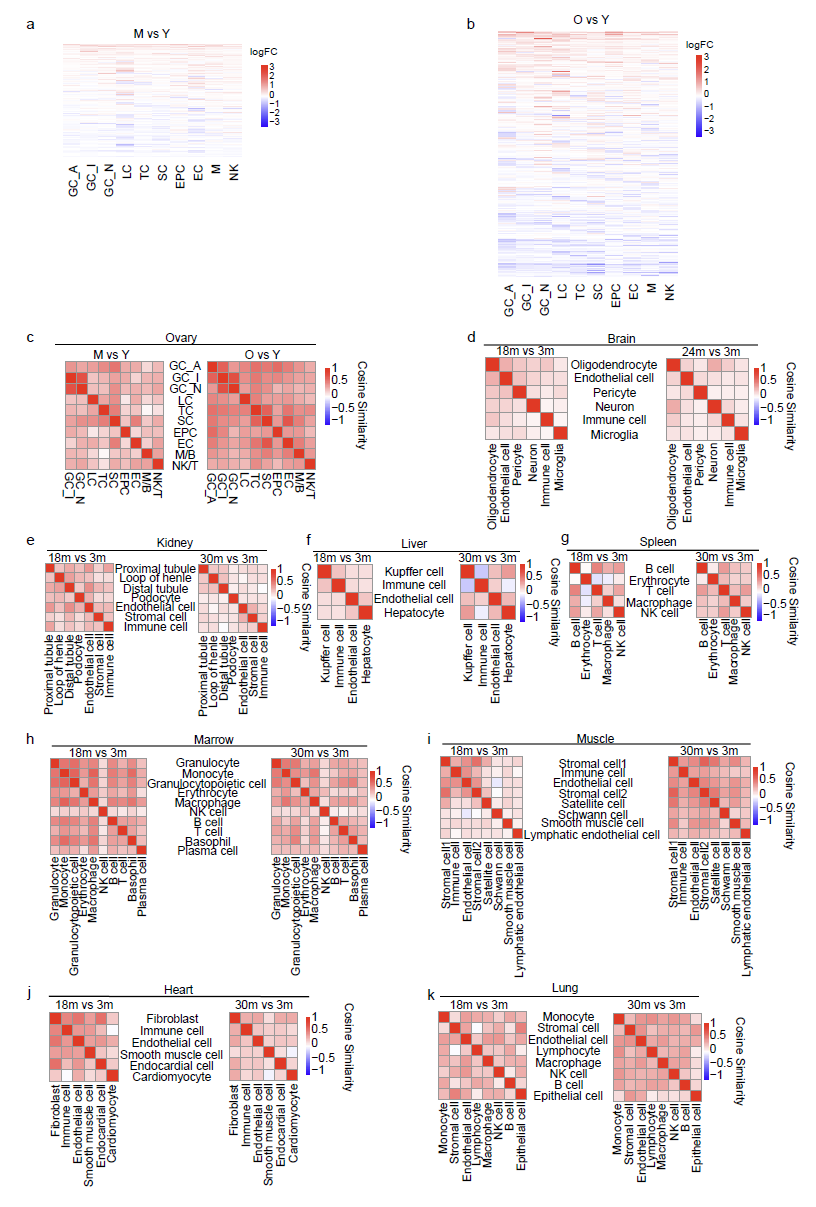

**Supplementary figure 2. Coordinated transcriptomic changes in mouse ovarian cells during aging. a-b,** Heat map showing log2 fold changes in gene expression of pairwise DEGs middle-aged ovary **(a,** M vs Y**)** and old ovary **(b,** O vs Y**)** comparing to young ovary across different cell types. **c-k,** Heat maps showing the pairwise cosine similarities of transcriptomic changes during aging between cell types in the mouse ovary **(c)** and brain **(d)**, kidney **(e)**, liver **(f)**, spleen **(g)**, marrow **(h)**, muscle **(i)**, and heart **(j)**, lung **(k)**.

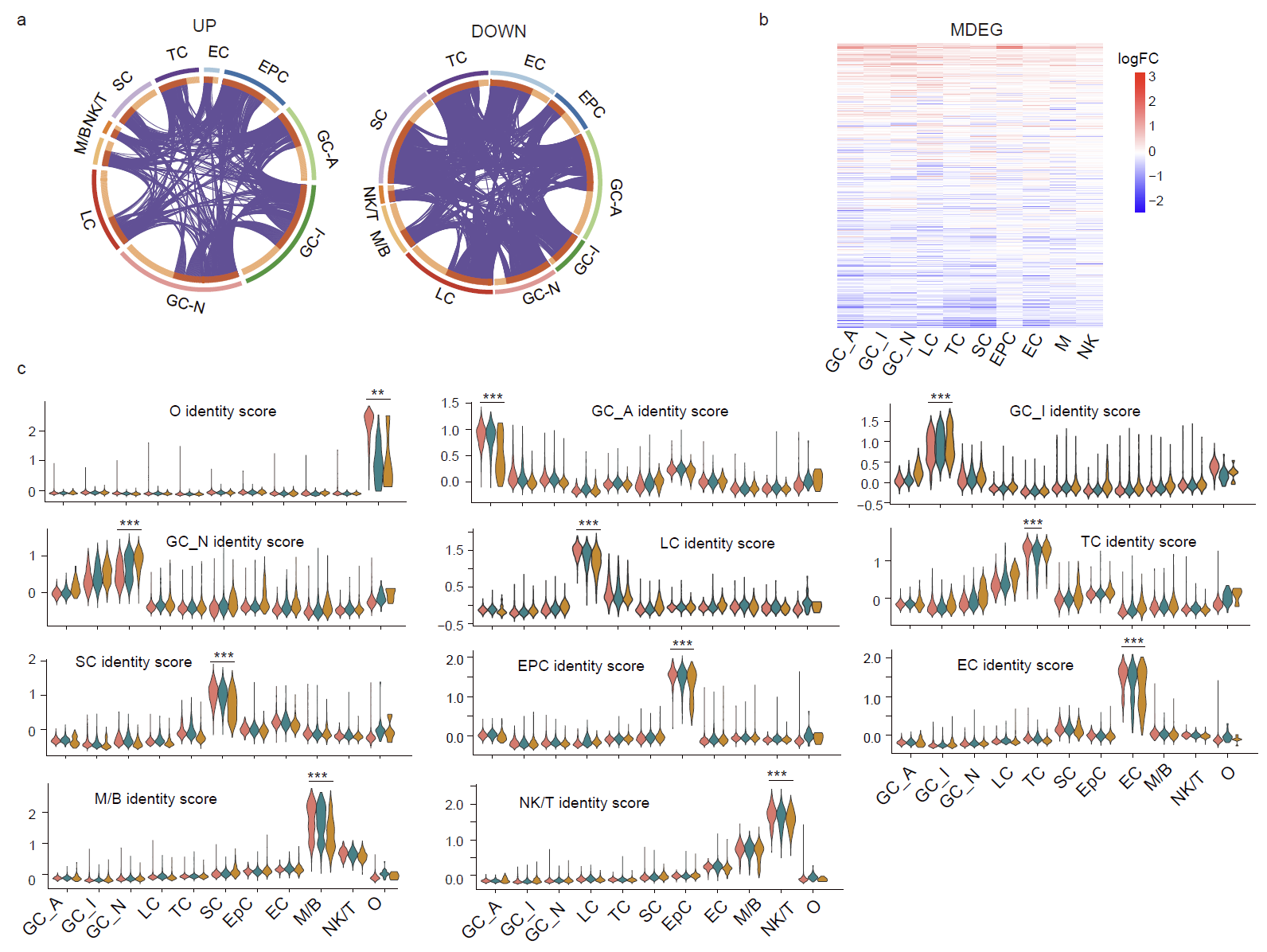

**Supplementary figure 3. Gene signatures of mouse ovarian aging. a,** Circos plots depicting the overlaps among gene lists of ovarian upregulated MDEGs (left) or downregulated MDEGs (right) for each cell type between old and young ovary. The inner circle represents gene lists, and purple curves link identical genes. The genes that hit multiple lists are colored in dark orange, and genes unique to a list are shown in light orange. **b,** Heat map showing log2 fold changes (O vs Y) in gene expression of MDEG during mouse ovarian aging in each cell type. **c,** Violin plots showing the cell identity score in each cell type at each age. (Wilcoxon test; **P<0.01, ***P<0.001).

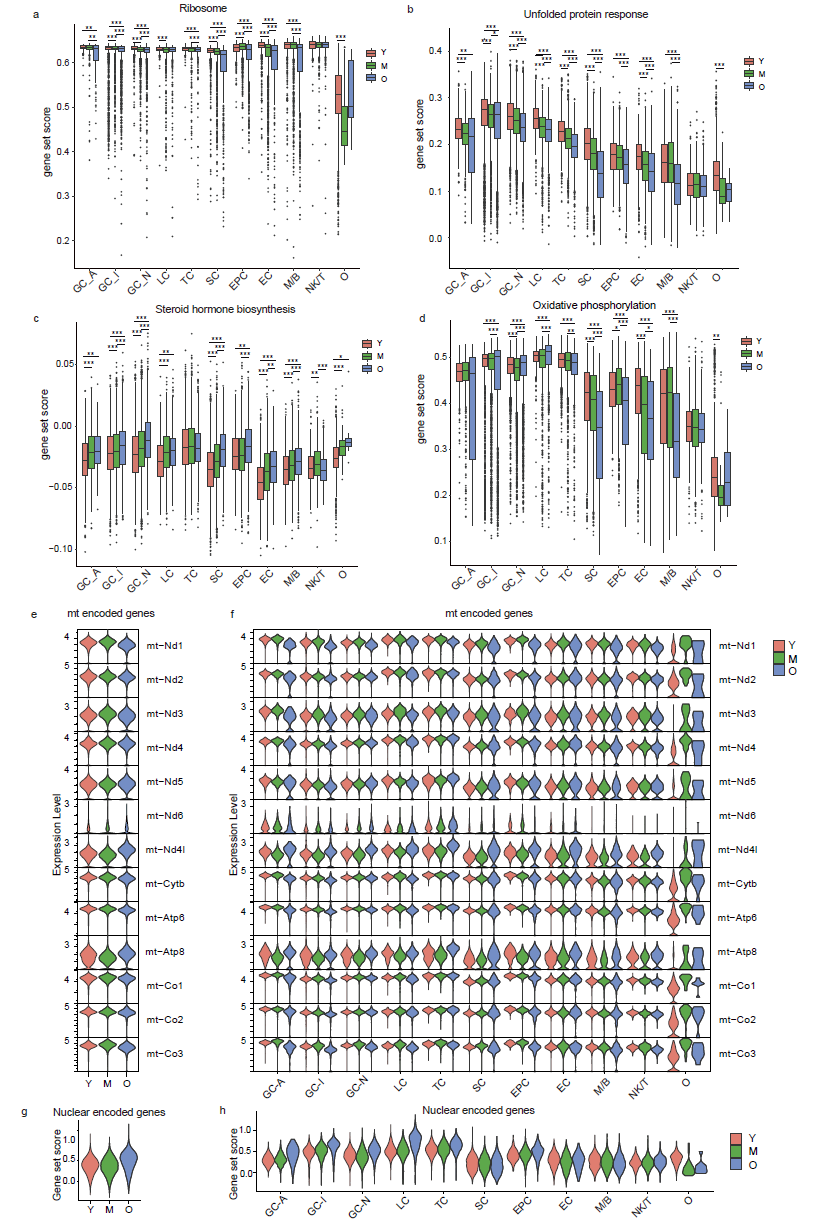

**Supplementary figure 4. Gene set scores of aging hallmarks related pathways in each cell type during mouse ovarian aging. a-d,** Box plots showing gene set scores of ribosome **(a)**, unfolded protein response **(b)**, steroid hormone biosynthesis **(c)** and oxidative phosphorylation **(d)** in each cell type at each age (Wilcoxon test; *P<0.05, **P<0.01, ***P<0.001). **e-f,** Violin plots showing expression level of mitochondria encoded genes of electron transport chain in all cell types **(e)** and in each cell type **(f)** at different ages. **g-h**, Violin plots showing the gene set score of nuclear encoded genes of electron transport chain in all cell types **(g)** and in each cell type **(h)** at different ages.

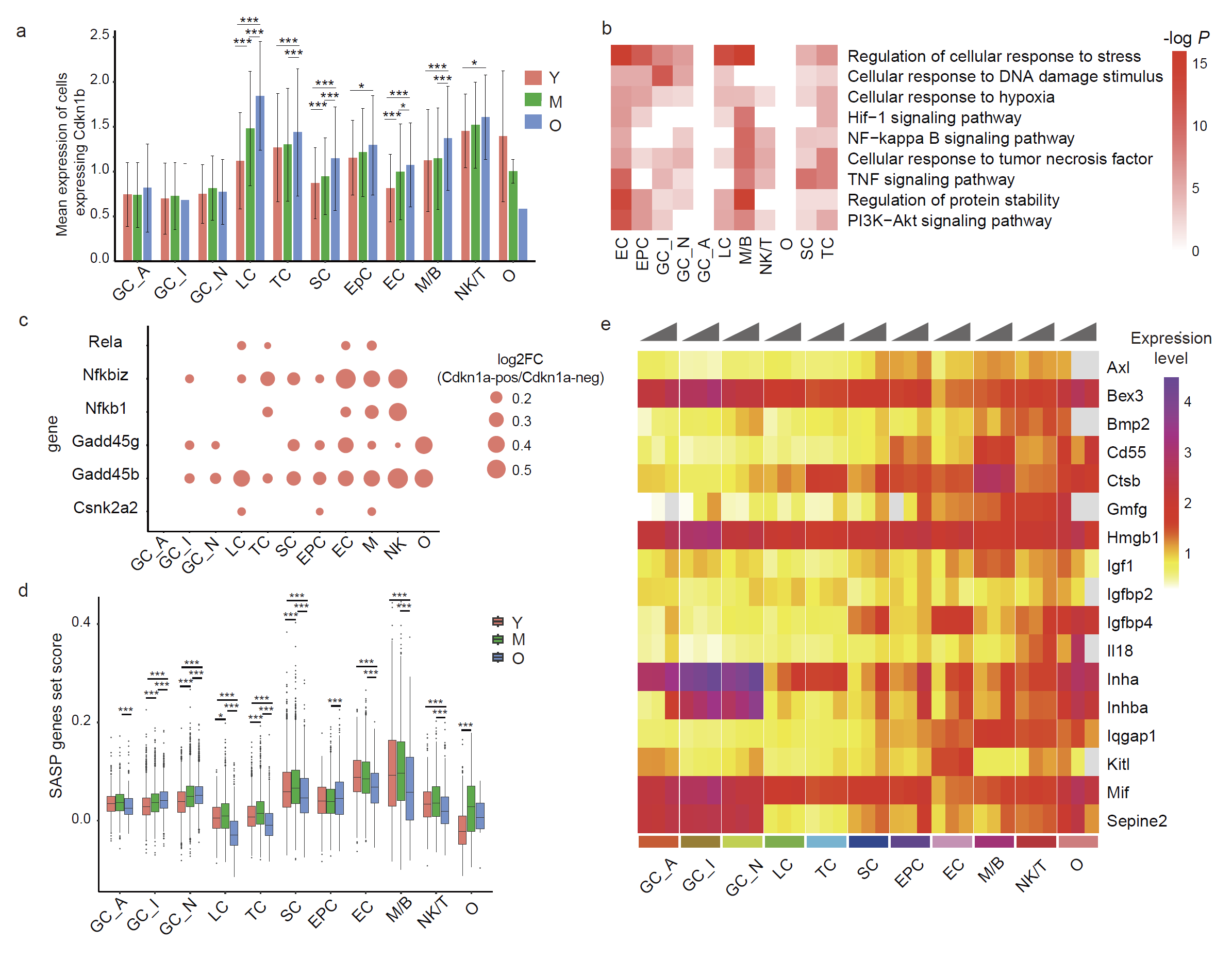

**Supplementary figure 5. Change of cellular senescence in each cell type during mouse ovarian aging. a,** Bar plots showing the expression of *Cdkn1b* in each cell type at each age. **b,** Heatmap showing representative GO terms of DEGs (MAST test, min.pct = 0.1, log FC > 0.1) between *Cdkn1a*+ cells and Cdkn1a- cells in each cell type. **c,** Dot plots showing the change in expression of representative genes in the NF-κB pathway in *Cdkn1a*^+^ cells compared to *Cdkn1a*^-^ cells. **d,** Box plots showing SASP gene set score in each cell type at each age group (Wilcoxon test; *P<0.05, **P<0.01, ***P<0.001). **e,** Heatmap showing the change of mean expression of upregulated monotonic DEGs overlapped with genes from SASP gene set.

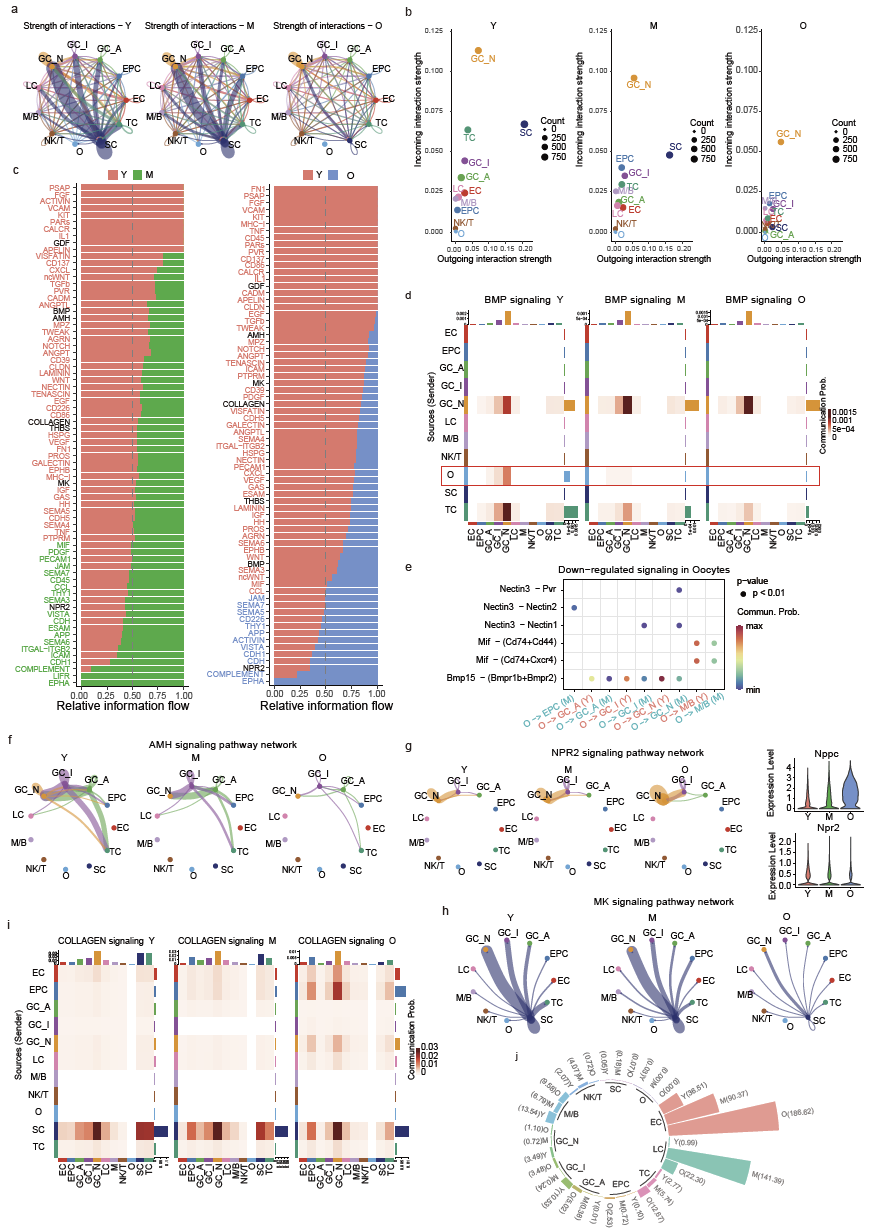

**Supplementary figure 6. Change of cellular interactions among different cell types during mouse ovarian aging. a,** Circle plots showing the strength of interactions at each age group. **b,** scatter plots showing cell types with significant changes in sending or receiving signals during mouse ovarian aging. **c,** stacked bar charts showing the significantly changed signaling information flow between middle-aged ovaries (left panel) or old ovaries (right panel) compared with young ovaries. The top signaling pathways colored red are enriched in young ovaries, and these colored greens are enriched in middle-aged or old ovaries. **d,** Heat map showing BMP signaling network in young, middle-aged, and old ovaries. Rows and columns represent sources and targets, respectively. Bar plots on the right and top represent the total outgoing and incoming interaction scores respectively. **e,** bubble plots showing the significantly downregulated signaling (ligand-receptor pairs) in oocytes during ovarian aging. **f,** Circle plots showing AMH signaling network in young, middle-aged, and old ovaries. **g,** Circle plots showing NPR2 signaling network in young, middle-aged, and old ovaries (left panel). Violin plots showing the expression of ligand and receptor of NPR2 signaling at each age group (right panel). **h,** Circle plots showing MK signaling network in young, middle-aged, and old ovaries. **i,** Heat map showing COLLAGEN signaling network in young, middle-aged, and old ovaries. **j,** ESAI_c of different ovarian cell types in young, middle-aged, and old groups.

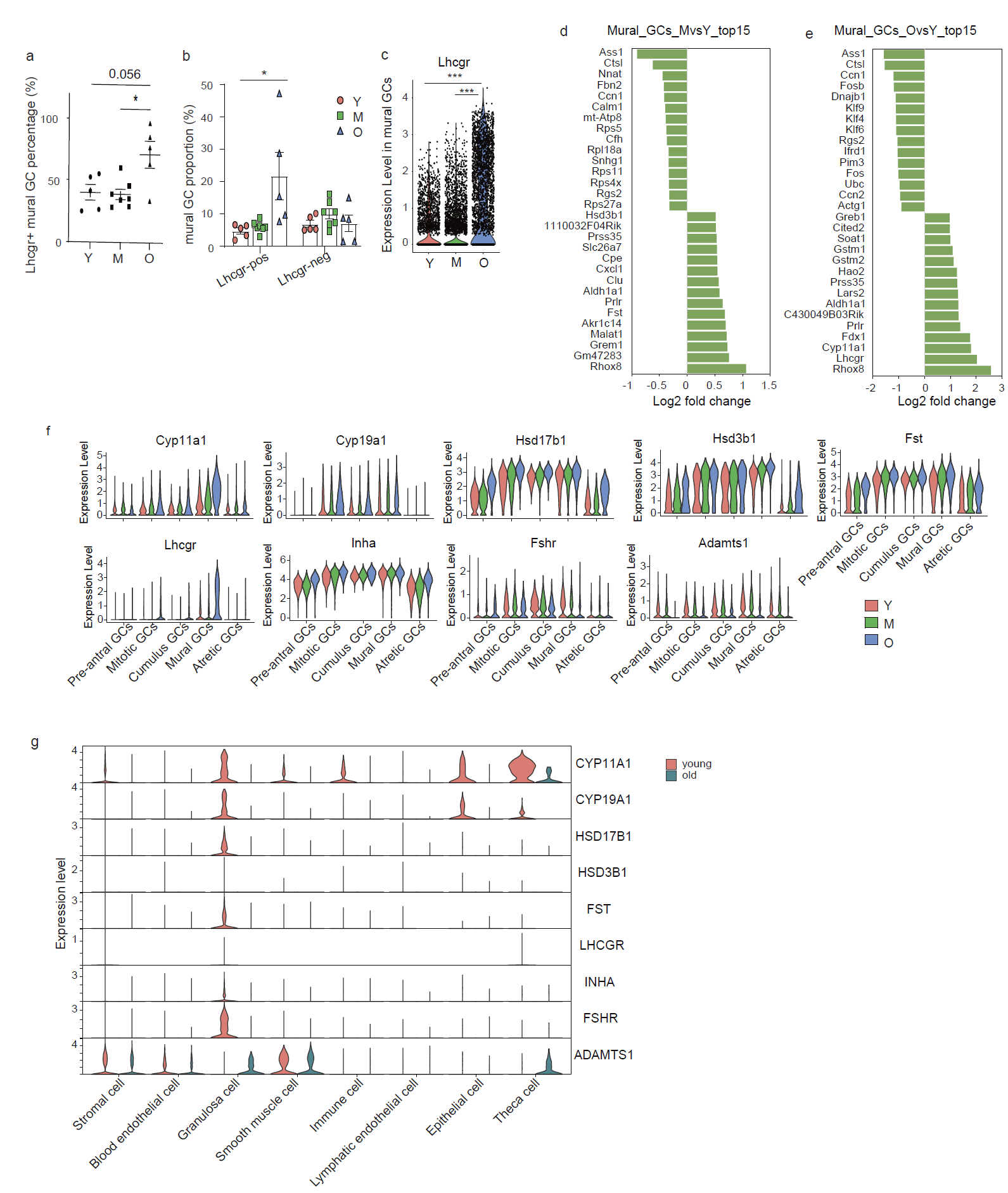

**Supplementary figure 7. Change of subpopulations of granulosa cells during mouse ovarian aging. a,** Scatter plots showing the percentage of Lhcgr^+^ cells in mural GCs. **b,** Bar plots showing the cell proportion of Lhcgr^+^ and Lhcgr^-^ mural GCs in young, middle-aged, and old ovaries. **c,** Violin plots showing the expression level of Lhcgr in mural GCs at each age group. **d-e,** Bar plots showing the top 15 upregulated and downregulated DEGs in mural GCs in middle-aged ovaries (**d**) and old ovaries (**e**) compared with young ovaries. **f,** Violin plots showing the expression of hormone-related genes in mouse granulosa cells at each age groups. **g,** Violin plots showing the expression of hormone-related genes in human ovarian cells in young and aged ovaries.

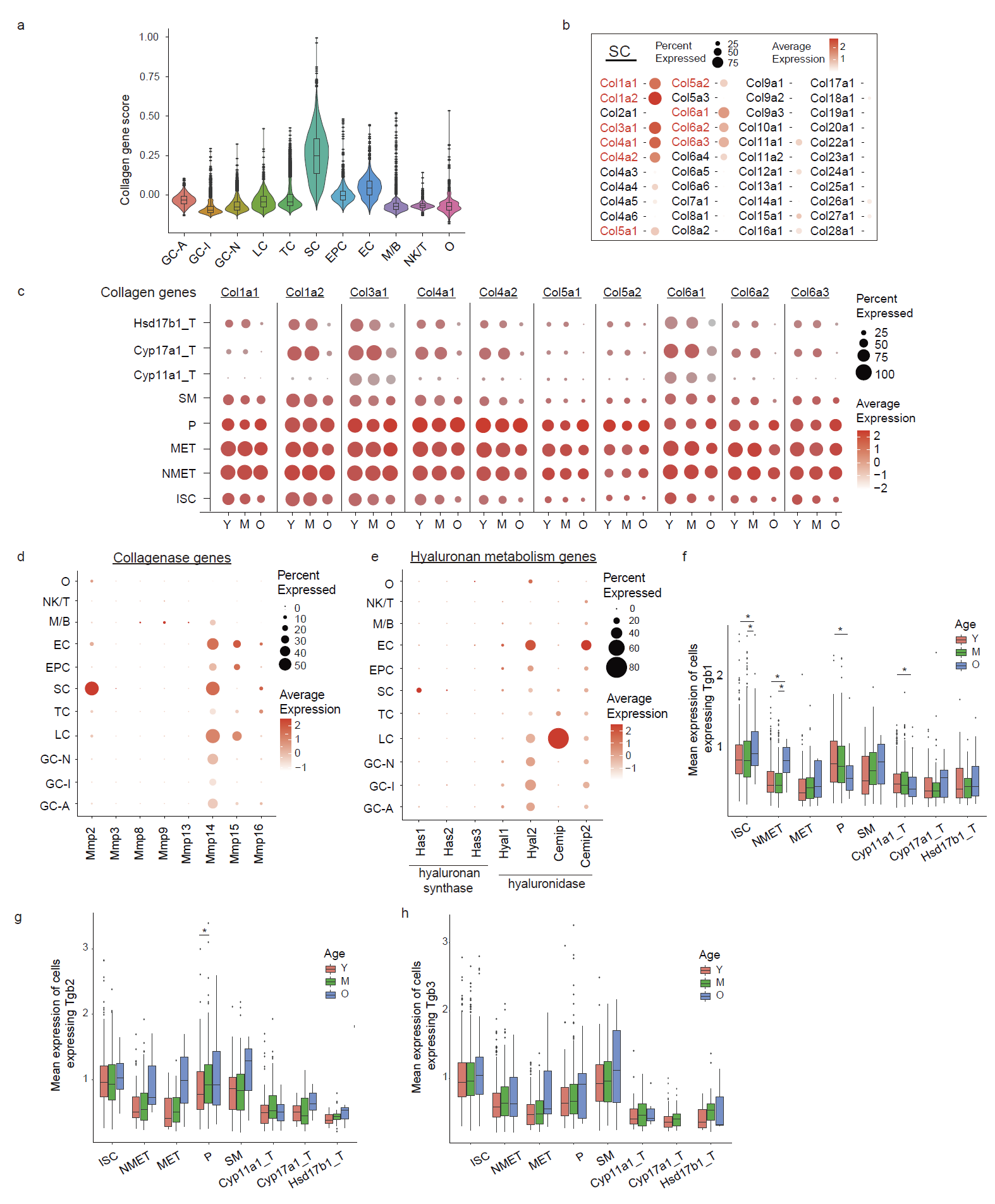

**Supplementary figure 8. Change of subpopulations of stromal and theca cells during mouse ovarian aging.** **a,** Violin and box plots showing the scores of collagen genes in each cell type of mouse ovarian cells. **b,** Dot plots showing the expression of each collagen gen in stromal cells. **c,** Dot plot showing the expression of relatively highly expressed collagen genes in each SC/TC subtype at each age. **d,** Dot plots showing the expression of collagenase genes in each cell type of mouse ovarian cells. **e,** Dot plots showing the expression of hyaluronan metabolism-related genes in each cell type of mouse ovarian cells. **f-h,** box plots showing the expression of Tgfb1 (**f**), Tgfb2 (**g**), and Tgfb3 (**h**) in each SC/TC subtype at each age. Wilcoxon test, *P<0.05.

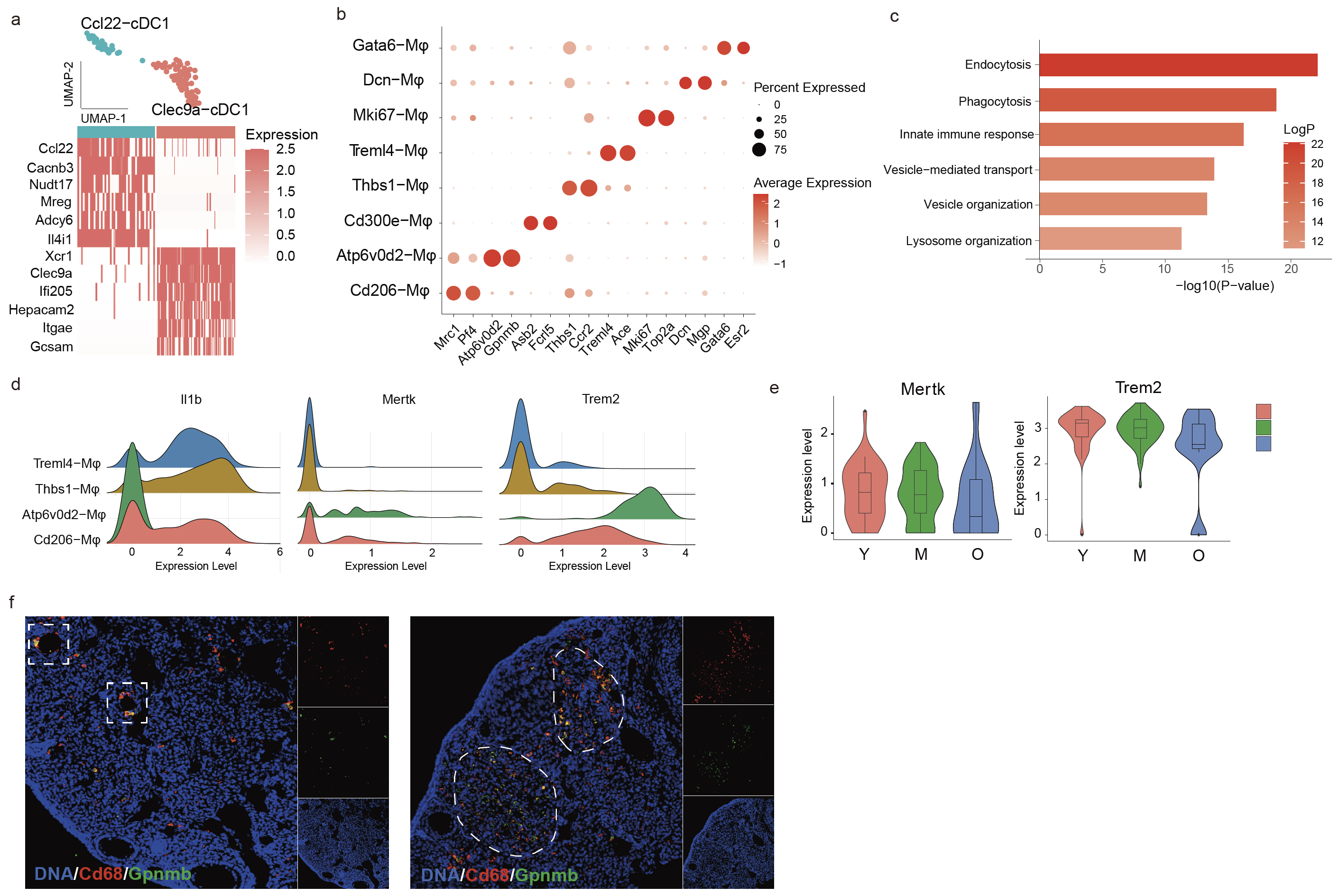

**Supplementary figure 9. Change of subpopulations of immune cells during mouse ovarian aging. a,** UMAP plots showing two cDC1 clusters (upper panel) and heatmap showing the expression of marker genes of each cluster (lower panel). **b,** Dot plots showing the expression of representative genes in subtypes of macrophage. **c**, GO analysis showing the functions that maker genes of Atp6v0d2- Mφ were enriched in. **d.** Ridge Plot showing the expression of Il1b, Mertk, and Trem2 in different macrophages. **e.** Violin plot showing the expression of Mertk and Trem2 in Atp6v0d2- Mφ at different age groups. **f**. Representative images of atretic follicles showing Gpnmb-positive macrophages in small degenerated atretic follicles (rectangle) and corpus luteum (circle) in young ovaries.
